# Evidence for divergent cortical organisation in Parkinson’s disease and Lewy Body Dementia

**DOI:** 10.1101/2025.06.25.661456

**Authors:** Angeliki Zarkali, Dr George Thomas, Ms Naomi Hannaway, Ms Ivelina Dobreva, Melissa Grant Peters, Martina F. Callaghan, Prof Mina Ryten, Rimona S Weil

## Abstract

Dementia is a defining feature of Lewy body disease: its timing and onset distinguish different clinical diagnoses, and its effect on quality of life is profound. However, it remains unclear whether processes leading to cognitive and motor symptoms in Lewy body disease differ. To clarify this, we used in-vivo neuroimaging to assess spatial gradients of inter-regional differences in structural and functional connectivity in 108 people across the Lewy body disease spectrum (46 Parkinson’s with normal cognition (PD-NC), 62 Lewy body dementia (LBD)) and 23 controls. We found divergent structural gradient differences with cognitive impairment: PD-NC showed increased inter-regional differentiation, whilst LBD showed overall gradient distribution similar to controls despite widespread organisational differences at the regional level. We then assessed cellular and molecular underpinnings of these organisational changes. We reveal similarities and also important differences in the drivers of cortical organisation between LBD and PD-NC, particularly in layer 4 excitatory neurons.

## Introduction

Lewy body diseases, characterised by the presence of alpha-synuclein aggregates forming Lewy bodies and Lewy neurites, are a group of heterogeneous clinical syndromes including Parkinson’s disease (PD), Parkinson’s disease dementia (PDD) and Dementia with Lewy bodies (DLB)^1^. Cognitive impairment in various degrees typically accompanies motor symptoms in Lewy body diseases and the timing of cognitive impairment defines disease subtypes, with PDD diagnosed when dementia occurs more than a year after the onset of motor symptoms and DLB if dementia occurs before^2,3^. PDD and DLB can also be grouped using the term Lewy body dementia (LBD) to refer to both conditions. The presence of dementia in addition to the motor symptoms characteristic of PD, is common, with up to 50% of patients with PD developing PDD^4^, and males and those with visuospatial impairment at higher risk^5^. Both PDD and DLB have a devastating impact on the quality of life of patients. Furthering our understanding of how PD with and without dementia differ is essential for disease characterisation and, ultimately, treatment.

Understanding how brain structure is altered in patients with and without dementia differ is particularly important. Several studies have shown changes in brain structure in association with cognitive decline in Lewy body diseases, including cortical atrophy^6,7^ and wide spread changes in grey^8–10^ and -white matter^11,12^ macrostructure. Although these studies provide useful insights, they were unable to take into account how different brain regions are interlinked at the cellular and molecular level across large-scale brain networks. This large-scale network architecture is underpinned by fundamental organisational gradients, which represent axes of continuous spatial transitions between brain regions^13^. Organisational gradients, have been shown in health to underly brain structural^14,15^ and functional connectivity^16^, accompany brain development^17,18^ and evolution^19,20^ and also align with the brain’s regional cytoarchitecture and gene expression^21,22^.

Differences in these regional cortical organisation gradients have also been found in psychiatric and neurological disease, with functional alterations described in both depression^23^ and schizophrenia^24,25^ and structural alterations in patients with epilepsy^26^. At the same time, these axes of organisation may be relevant to cognition, with structural gradient expansion and higher inter-regional differentiation between unimodal and more transmodal regions in adolescence supporting executive function^27^. However, they have not yet been examined in cognitive disorders such as LBD.

We have previously shown that differences in structure-function coupling in patients with PD at higher risk of dementia follow these established gradients of cortical organisation^28^. However, no imaging studies have yet examined whether cortical reorganisation accompanies cognitive decline in Lewy body diseases nor the spatial pattern of these changes. Recent evidence from post-mortem gene expression, suggests that cortical organisation is disrupted in both PD and LBD but in divergent ways^29^: intra-individual regional analysis of single-cell RNA sequencing from two cortical regions with different alpha-synuclein burden showed increased inter-individual differences in PD but attenuation of regional identity in PDD, whilst there was also evidence of distinct cellular and molecular underpinnings of this reorganisation^29^. Whether inter-regional differentiation is altered beyond these two regions at a whole brain level and how such whole-brain alterations might differ between PD with intact cognition (PD-NC) and LBD has yet to be established.

We aimed to examine in-vivo large-scale differences in cortical structural and functional organisation between patients with LBD, PD-NC and unaffected controls (HC) (*Figure 1*). To do this we compared both the overall distribution and vertex-wise differences in in-vivo structural and functional gradients, derived from diffusion-weighted and resting-state functional MRI (rsfMRI) respectively between LBD, PD-NC and HC. We found increased widening of structural gradients, therefore increased inter-regional differentiation in PD-NC and apparent normalisation of inter-regional differentiation in LBD compared to controls.

**Figure 1.**
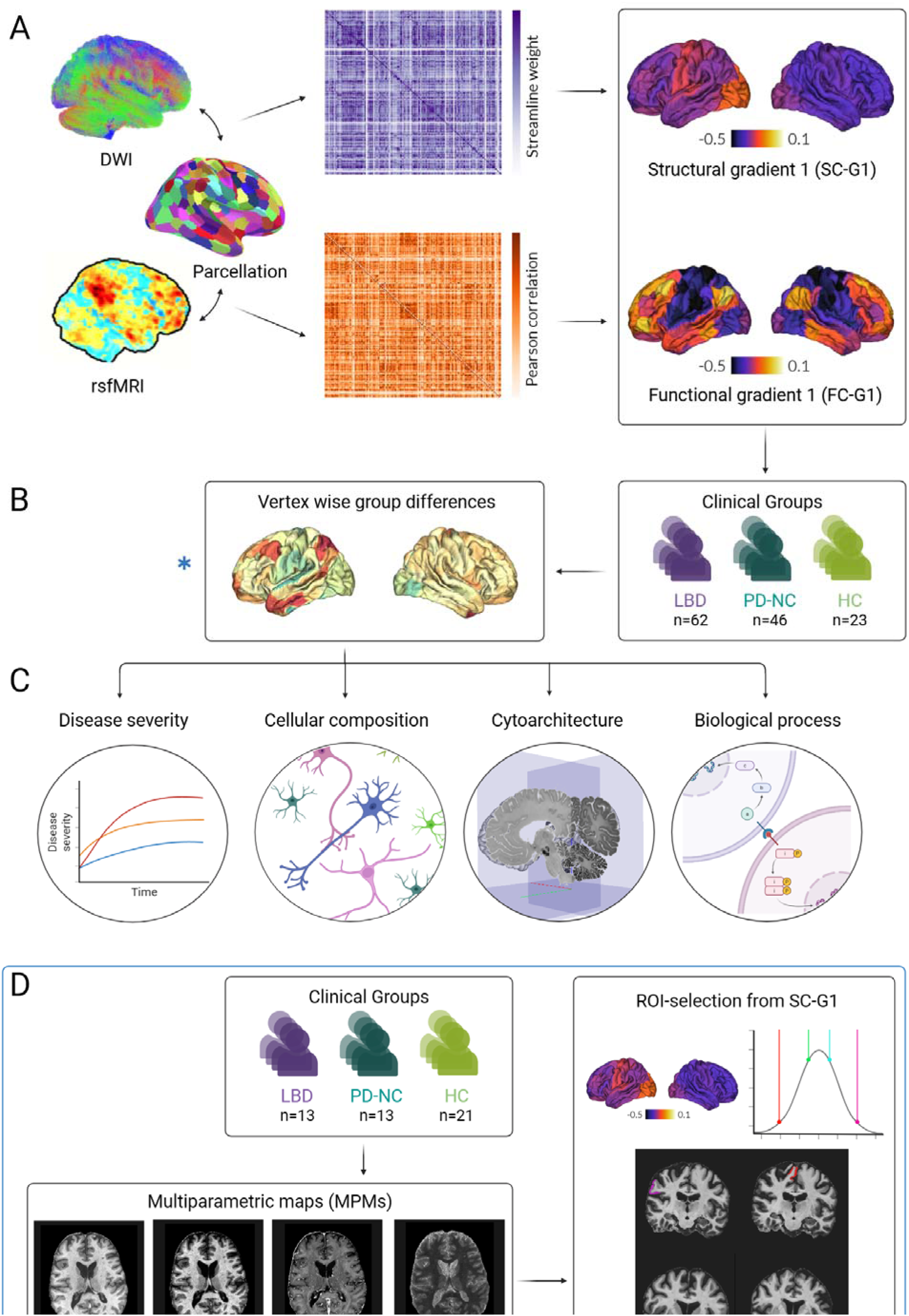
Assessing cortical organisation across the spectrum of Lewy body disease. **A. Individualized structural and functional gradient construction**. Diffusion-weighted (DWI) and resting state functional MRI (rsfMRI) images were obtained from each participant. Structural T1-weighted imaging was used to parcellate the brain into 200 cortical regions of interest using the Schaeffer parcellation^82^. The same parcellation was used to derive structural connectivity matrices weighted by streamlines between any two regions (purple) and functional connectivity matrices, weighted by Pearson correlation between any two regions (orange) for each participant. We then applied diffusion map embedding, a non-linear dimensionality reduction technique^16,83^ to identify gradients (eigenvectors) of the main spatial axes in inter-regional similarity of structural and functional connectivity. The average principal gradient for structural (SC-G1) and functional connectivity (FC-G1) in healthy controls is shown. **B. Surface-based linear models revealed significant differences in structural gradient scores (SC-G1) between groups.** Gradients were compared between 3 groups: 62 patients with Lewy body dementia (LBD), 46 patients with Parkinson’s disease with intact cognition (PD-NC) and 23 unaffected controls (HC). We used surface-based linear models controlling for age and sex to compare gradient scores between LBD vs HC and PD-NC vs HC. We validated changes in structural organisation using a region-of-interest analysis (in regions across the SC-G1 spectrum) and quantitative ultra-high field 7 Tesla MRI in a separate cohort of patients. **C. Neural contextualisation of cortical structural organisation alterations**. We assessed whether the 1) the structural SC-G1 gradient alterations seen in Lewy body disease were related to cognitive and motor disease severity and whether 2) the spatial distribution of vertex-wise SC-G1 alterations seen in Lewy body disease (LBD vs HC and PD vs HC) was correlated with the spatial distribution of specific neuronal and glial cell types, cytoarchitectural axes of organisation, and expression of genes linked to specific biological processes and pathways. **D. Replication analysis in a separate cohort using 7 Tesla quantitative MRI**. Multiparametric maps (MPMs) were acquired at 7 Tesla for each participant, including maps sensitive to myeline such as longitudinal relaxation rate (R1) and magnetisation transfer saturation (MTsat), iron-sensitive map effective transverse relaxation rate (R2*), and map sensitive to overall water content, proton density. We then extracted mean MPM signal for four regions of interest (ROIs) of the same Schaeffer parcellation used in our main analysis. ROIs were selected based on their SC-G1 ratings from the main 3T cohort and whether they differed in LBD compared to controls. Thus, two regions were selected from the extremes of the gradient distribution which differed between LBD and controls (“RH_SalVentAttn_TempOccPar_3”, and “RH_SomMot_18”) and two were selected from the middle of the gradient distribution and which did not show differences between LBD vs controls (“RH_Default_Temp_1”, “LH_Default_Temp_1”). We assessed for ROI*Group interaction using mixed linear models with age and sex as covariates to assess whether interregional differences in MPM signal differed significantly between the three clinical groups.

However, there were widespread differences in regional gradient rankings in LBD relative to controls for the primary structural gradient (SC-G1) suggesting significant local and regional re-organisation despite normalisation of the overall distribution. We validated these differences using region-of-interest (ROI) analysis and 7T quantitative MRI (qMRI) in a separate cohort. We confirmed that inter-regional differences in mean qMRI values for myelin sensitive metrics across regions of different SC-G1 rankings, significantly differed between groups. Then, we examined whether these differences in cortical structural organisation were linked to disease severity, by correlating the overall difference from control SC-G1 ratings, with cognitive and motor severity measures. We show structural organisation was behaviourally relevant in LBD and specific to cognitive severity. Finally, we aimed to understand whether disease-related gradient alterations were associated with normative topographical variations in excitatory and inhibitory neurons, cortical cytoarchitecture and global gene expression. We reveal similarities but also important differences in the drivers of changes in cortical organisation between LBD and PD-NC, particularly in excitatory neurons.

## Results

The overall study methodology is seen in *Figure 1*. 62 LBD patients (including DLB, PDD and PD with mild cognitive impairment (PD-MCI)), 46 PD patients with stable cognition (PD-NC) and 23 healthy age-matched controls (HC) were included. Demographics and clinical assessments are seen in *Table 1*. Full details of cognitive assessments and characteristics of different LBD subgroups are seen in *Supplementary Tables 1 and 2*.

**Table 1.** Demographics and clinical characteristics.

| Characteristic | Controls (HC)<br>n=23 | Parkinson's with<br>normal<br>cognition (PD-NC)<br>n=46 | Lewy Body<br>Dementias<br>(LBD)<br>n=62 | p-value |
| --- | --- | --- | --- | --- |
| **Age** | 66.8 (8.6) | 62.3 (7.2) | 70.1 (6.7) | **p=0.030<sup>c</sup>** |
| **Male (%)** | 11 (47.8) | 22 (47.8) | 50 (80.6) | **p<0.001<sup>a,c</sup>** |
| **Right-handed (%)** | 21 (91.3) | 45 (97.8) | 53 (85.5) | p=0.190 |
| **Years of education** | 17.6 (2.4) | 17.0 (2.6) | 16.3 (3.7) | p=0.163 |
| **MOCA** | 28.6 (2.6) | 28.9 (1.2) | 23.4 (5.4) | **p<0.001<sup>a,c</sup>** |
| **MMSE** | 28.9 (1.9) | 29.2 (1.2) | 26.6 (3.5) | **p<0.001<sup>a,c</sup>** |
| **HADS anxiety** | 3.8 (3.7) | 4.6 (2.7) | 6.6 (4.7) | p=0.012<sup>a</sup> |
| **HADS depression** | 2.6 (2.9) | 3.5 (2.4) | 6.2 (3.1) | **p<0.001<sup>c</sup>** |
| **Composite cognitive score** | - | 0.1 (0.5) | -1.8 (1.7) | **p<0.001<sup>a,c</sup>** |
| **Years of diagnosis** | - | 4.7 (2.9) | 4.0 (4.0) | p=0.090 |
| **UPDRS total** | - | 49.9 (15.5) | 66.3 (28.5) | **p=0.008** |
| **UPDRS part 3** | - | 28.3 (8.3) | 32.8 (15.6) | p=0.307 |
| **Timed Up and GO** | - | 8.9 (1.9) | 10.3 (4.5) | p=1.000 |
| **Functional Assessments Questionnaire** | - | 1.7 (1.9) | 8.3 (6.4) | **p<0.001** |
| **CAF** | - | 2.4 (3.1) | 4.7 (3.7) | p=0.171 |
| **One Day Fluctuations Questionnaire** | - | 1.4 (2.0) | 6.4 (6.9) | **p=0.020** |
| **LEDD** | - | 500.0 (389.7) | 515.9 (253.8) | p=3.719 |
| **UMPDHQ** | - | 0.8 (1.9) | 2.9 (3.3) | **p<0.001** |
| **RBDSQ** | - | 4.5 (2.8) | 7.0 (4.0) | **p=0.012** |

<sup>a</sup>: difference between LBD-HC, <sup>b</sup>: difference between PD-HC, <sup>c</sup>: difference between LBD-PD. In bold significant differences between LBD and PD. All results shown as mean (standard deviation) except otherwise indicated.
CAF: Clinician Assessment of Fluctuations; HADS: Hospital Anxiety and Depression Scale, LEDD: Levodopa Equivalent Daily Dose; MMSE: Mini-Mental State Examination; MOCA: Montreal Cognitive assessment; RBDSQ: REM Sleep Behaviour Disorder Sleep Questionnaire; UMPDHQ: University of Miami PD

### Structural connectivity gradients are distinct in LBD and PD-NC

First, we derived cortex-wide structural and functional connectivity gradients for each participant. To assess how inter-regional gradients differ in LBD and PD-NC, we then estimated eigenvectors describing spatial gradients of systematic cortical variation in brain structural and functional connectivity also known as cortical gradients. We compared structural and functional gradients between controls, PD-NC and LBD. First, we assessed the overall gradient distribution for the primary structural connectivity gradient (SC-G1) and the second structural connectivity gradient (SC-G2): we found that PD-NC patients showed expansion (scores shifting further away from the midpoint) of structural connectivity gradients compared to controls and LBD (SC-G1: Kruskal-Wallis H=7.87, p=0.019; SC-G2: H=17.0, p<0.001). This reflects that regional alterations in structural connectivity in PD-NC result in increased differentiation between different brain regions. In contrast, LBD patients showed constriction of the overall gradient distribution compared to PD-NC with a slight qualitative reduction of regions in the extreme of gradients compared to controls (SC-G1: *Figure 2A; SC-G2: Supplementary* Figure 1). This suggests that increased inter-regional differentiation is a feature of PD-NC but not LBD.

**Figure 2.**
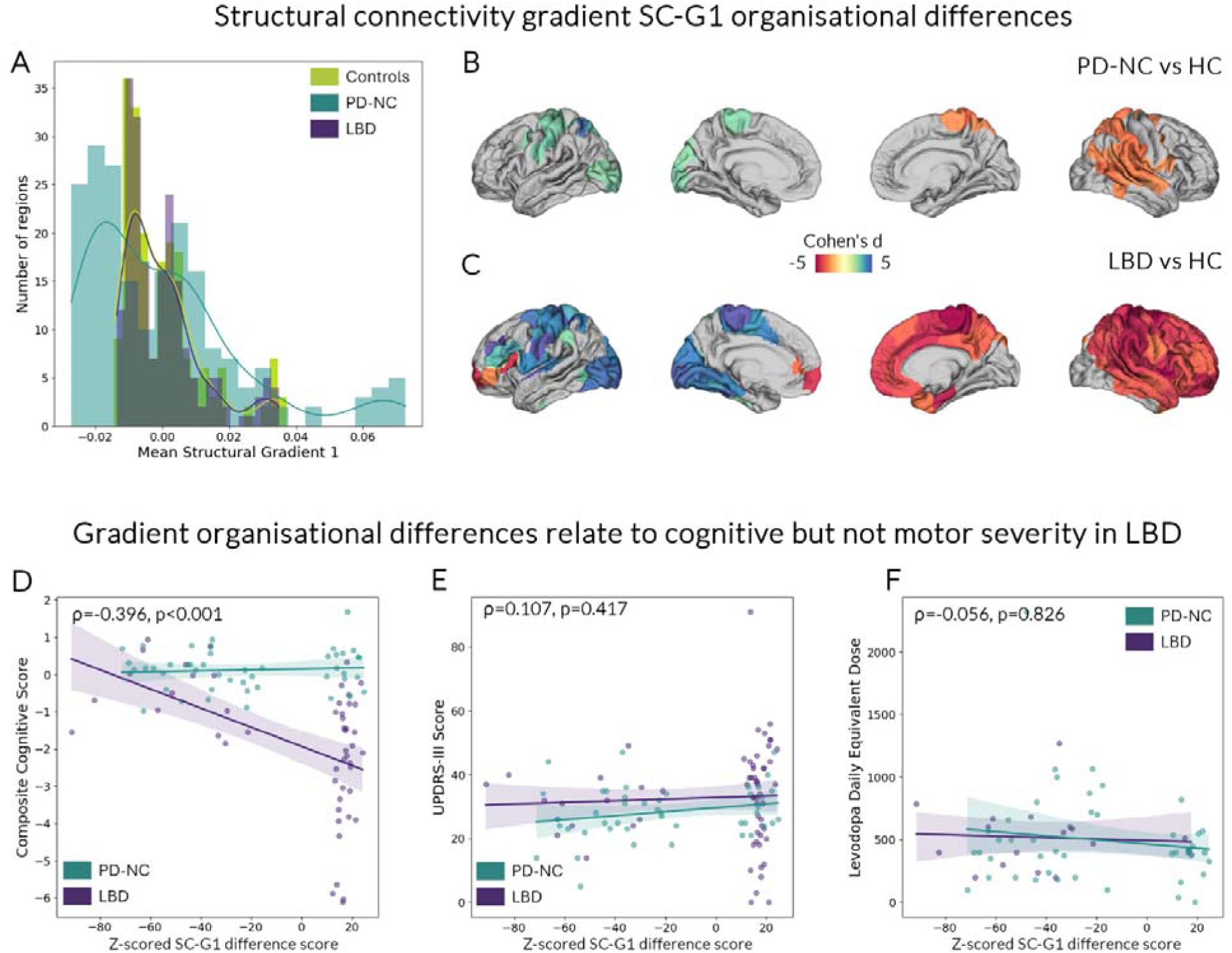
Structural connectivity gradient alterations are seen in Lewy body disease and are specific to cognitive severity. **Overall gradient distribution in the principal structural gradient (SC-G1) differed between controls (HC), Parkinson’s patients with normal cognition (PD-NC) and patients with Lewy Body Dementias (LBD) (A)** with expansion of gradient scores in PD-NC reflecting increased inter-regional differentiation in structural connectivity (Kruskal Wallis H=7.87, p=0.019). **Regional differences in structural gradient scores were seen in both PD-NC (B) and LBD patients compared to HC (C).** Surface-based linear models controlling for age and sex revealed significant differences in SC-G1 gradient scores between groups. Only statistically significant clusters after multiple comparisons correction (p_FWE_ <0.05) are shown. Colour scale depicts effect size per vertex (blue colours increases in gradient scores; red colours reductions in gradient scores). **Gradient reorganisation is specific to cognitive severity in LBD.** For each participant, a composite gradient change score was calculated from the Z-scored gradient value for each region compared to HC gradient values for that region, summed across the 200 regions. Composite gradient scores were correlated with composite cognitive scores (**D**) but not motor scores (UPDRS-III: Unified Parkinson’s Disease Rating Scale Part 3) (**E**) or levodopa daily equivalent doses (**F**).

Next, to better understand regional contribution to global cortical organisation and how a given region’s position within the cortical hierarchy may differ in Lewy body diseases, we used surface-based mixed linear models (age and sex as covariates, FWE-correction for multiple comparisons, cluster threshold 0.01), with each vertex allocated to its corresponding structural gradient value. Differences in gradient values in PD-NC compared to controls were concentrated in the extremes of the gradient distribution for both gradients: for SC-G1, there were changes in scores primarily in unimodal regions such as the sensorimotor and left visual cortex (*Figure 2B*). For SC-G2 there were reductions in gradient values of left prefrontal regions (*Supplementary* Figure 1). In contrast, and despite a similarity of the overall gradient distribution in LBD and control groups, there were bidirectional, bilateral and widespread differences in their spatial organisation in LBD compared to controls for both structural connectivity gradients SC-G1 (*Figure 2C*) and SC-G2 (*Supplementary* Figure 1).

When directly comparing PD-NC and LBD at the vertex rather than region-of-interest (ROI) level, we found no statistically significant differences (uncorrected results are presented in *Supplementary* Figure 2). In contrast to the pronounced differences in structural connectivity gradients shown in our cohort, functional connectivity gradients did not differ between any groups.

Several replication analyses (with structural connectivity gradients derived using different sparsity thresholds) demonstrated the robustness of structural gradient differences in LBD and PD-NC compared to controls (presented in *Supplementary* Figure 3).

### Differences in inter-regional differentiation are replicated using ultra-high field quantitative MRI in a different cohort of LBD, PD-NC and controls

To ensure the robustness of our results, we replicated our findings in a separate cohort of LBD, PD-NC and controls using different MR acquisitions and analyses. We used 7 Tesla quantitative MRI (7T qMRI) to test qMRI values from regions at the extremes of SG-G1 distribution (that differed between groups in our main analysis) against regions from the middle of the distribution (which did not differ between groups).

We included 13 LBD, 13 PD-NC and 21 age-matched controls in our replication 7T cohort. Demographics and results of clinical assessments in the replication cohort are presented in *Supplementary Table 3*. We computed quantitative multiparametric maps (MPM) including proton density, longitudinal relaxation rate (R1), effective transverse relaxation rate (R2*), and magnetisation transfer saturation (MTsat)). We extracted mean MPM values for four ROIs of the Schaeffer parcellation based on their SC-G1 ratings from the main 3T cohort and whether they differed in LBD compared to controls. Thus, two regions were selected from the extremes of the gradient distribution which differed between LBD and controls (“RH_SalVentAttn_TempOccPar_3”, and “RH_SomMot_18”) and two were selected from the middle of the gradient distribution and which did not show differences between LBD vs controls (“RH_Default_Temp_1”, “LH_Default_Temp_1”). We then used mixed linear models accounting for age and sex to study the ROI*Group (LBD, PD-NC, HC) interaction and assess whether interregional differences in MPM values differed significantly between groups.

We found a significant ROI*Group interaction for myelin sensitive maps, both R1 (p=0.045) and MTsat (p=0.022) but not for proton density or R2*. For MTsat, the group difference in inter-regional MTsat values was driven by the LBD group in the “RH_SomMot_18” region (β=-0.131, p=0.003). For R1 there were no significant regions or groups driving the overall ROI*Group interaction in pairwise comparisons, suggesting the overall difference was not driven by a single region. Full results of the replication analysis are presented in *Supplementary* Figure 4.

### Structural connectivity organisation changes are behaviourally relevant and specific to cognitive performance and Lewy body pathology

Next, we examined whether the changes in structural cortical organisation in Lewy body disease were related to disease severity. To do this we calculated for each participant, a composite gradient difference score, derived from the Z-scored SC-G1 gradient value for each region compared to the control gradient values for that region, then summed across the 200 regions. We correlated composite gradient difference scores with measures of disease severity (cognitive severity: Montreal Cognitive Assessment (MOCA) and mini-mental state assessment (MMSE), and a composite cognitive score combining detailed cognitive assessments across 5 cognitive domains; and motor severity: Movement Disorders Society Unified Parkinson’s Disease Scale part III (MDS-UPDRS-III) and “timed up and go” (TUG) score) within each disease group (LBD and PD-NC). We found that more pronounced deviations from normative cortical organisation (higher composite gradient difference scores) were associated with lower cognitive performance across all cognitive measures within LBD patients (SC-G1: MOCA: Spearman ρ=-0.292, p=0.021; MMSE: ρ=-0.244, p=0.05; composite cognitive score: ρ=-0.335, p=0.017) but not for PD-NC (all non-significant; *Figure 2D*).

In contrast there was no correlation between composite gradient difference scores and motor measures (MDS-UPDRS-III or TUG scores, *Figure 2E*) or levodopa equivalent daily dose (LEDD) (*Figure 2F*). These findings suggest that the differences in structural cortical organisation seen in LBD and PD-NC patients are specific to cognitive decline in Lewy body diseases, rather than other clinical features such as motor severity.

### Neuronal underpinnings of structural gradient differences show divergent contributions in PD-NC and LBD, with excitatory neurons affected only in LBD

We then aimed to evaluate whether structural gradient differences are associated with regional variations in specific neuronal and glial cell populations. To do this, we correlated the unthresholded t-map of gradient differences in LBD vs controls and PD-NC vs controls with regional gene expression of cell-specific gene markers (against 1000 spatially correlated spin permutations). We found that SC-G1 differences in LBD compared to controls were associated with the regional distribution of excitatory (ρ=0.192, p_spin_= 0.039) and inhibitory neuronal cells (ρ=-0.417, p_spin_<0.001), as well as oligodendrocytes (ρ=-0.193, p_spin_=0.021). We further examined whether specific cell types were driving the correlation with excitatory or inhibitory neuronal cell populations. The relationship with excitatory cell distribution was driven by RORB expressing layer 4 neurons (ρ=0.257, p_spin_= 0.022), with no other excitatory neuronal marker showing significant correlation. This suggests that regions differentially expressing inhibitory and excitatory cell populations are more likely to show changes in cortical hierarchical organisation in LBD; with regions rich in excitatory and poor in inhibitory cells (higher E-I ratio, particularly in RORB layer 4 neurons) showing increased SC-G1 values and regions with lower E-I ratio and poorer in oligodendrocytes showing contraction in SC-G1 values (*Figure 3A*).

**Figure 3.**
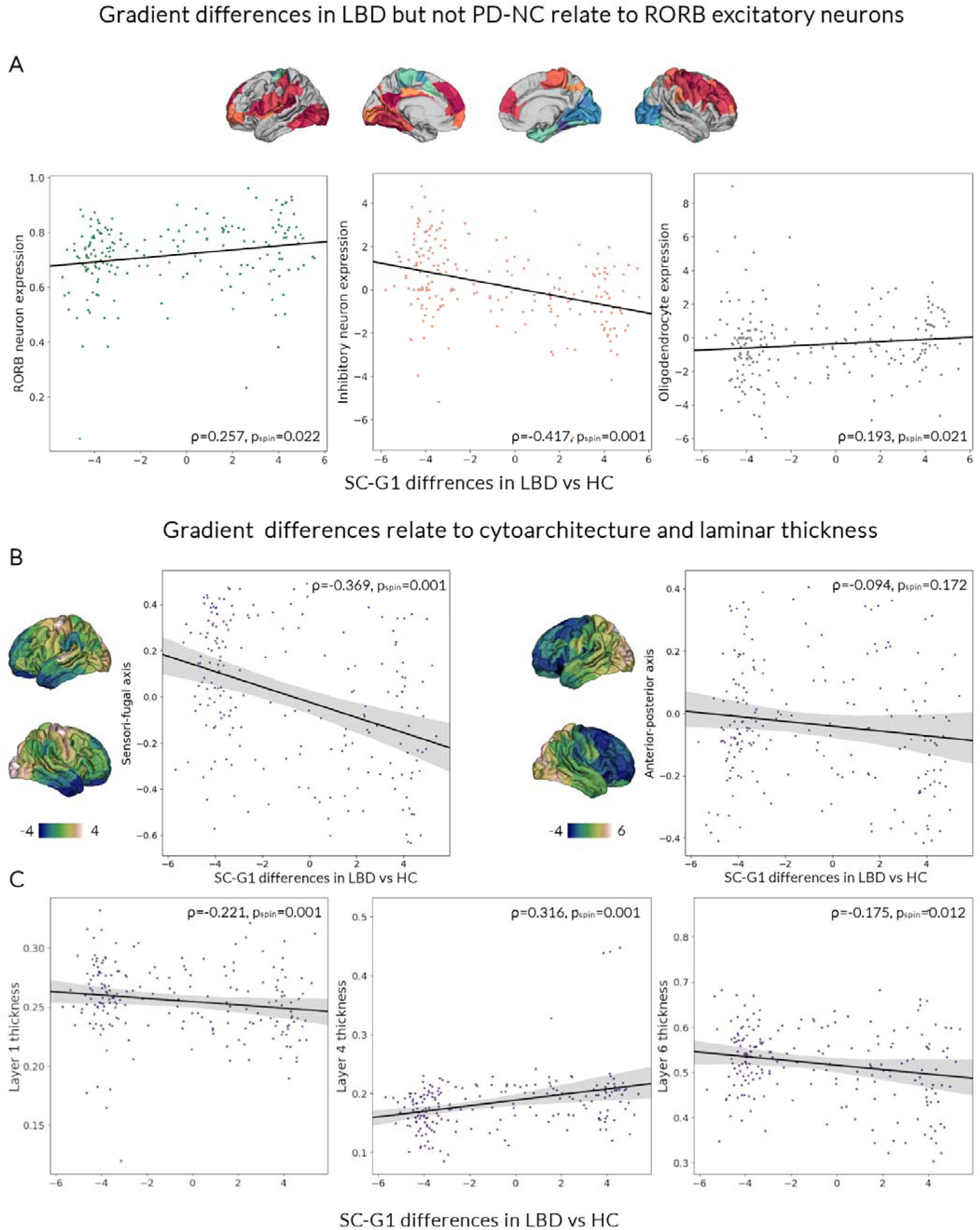
Neuronal underpinnings of the structural gradient alterations in LBD patients. A. The spatial distribution of structural gradient alterations in Lewy body dementia participants (LBD) is significantly correlated with the spatial distribution of normative gene expression markers of RORB excitatory neurons, inhibitory neurons and oligodendrocytes. The unthresholded t-map of primary structural gradient (SC-G1) changes in LBD vs healthy controls (HC) was correlated with normative regional gene expression of cell-specific gene markers (against 1000 spatially correlated spin permutations) from the Allen Human Brain Atlas^31^. SC-G1 changes in LBD compared to HC were positively associated with the regional distribution of RORB expressing layer 4 neurons (left) and overall inhibitory neuronal cells (middle) as well as oligodendrocytes (right). B. Structural gradient alterations in LBD follow a sensori-fugal axis of cytoarchitecture. The SC-G1 t-map of changes in LBD vs HC) was correlated to atlases of normative cytoarchitectural organisation from BigBrain^93^ (against 1000 spatially correlated spin permutations). SC-G1 alterations were spatially correlated with the sensory-fugal but not anterior-posterior axis of cytoarchitectural differentiation. C. Structural gradient alterations in LBD show specific laminar contributions SC-G1 gradient change t-maps (LBD vs HC) were negatively correlated with the regional variation of layer 1 (left) and layer 6 (right) cortical thickness and positively correlation with layer 4 cortical thickness (middle).

In contrast, regional changes in gradients in PD-NC compared with controls were only correlated with inhibitory but not excitatory neuronal distribution (ρ=-0.376, p_spin_= 0.004), and with regional distribution of oligodendrocytes (ρ=0.199, p_spin_= 0.018); with regions richer in excitatory neurons and poorer in oligodendrocytes more likely to show contraction in SC-G1 values in PD-NC patients. Full results of neuronal underpinnings of structural gradient alterations in LBD and PD-NC are shown in *Table 2*.

**Table 2.**
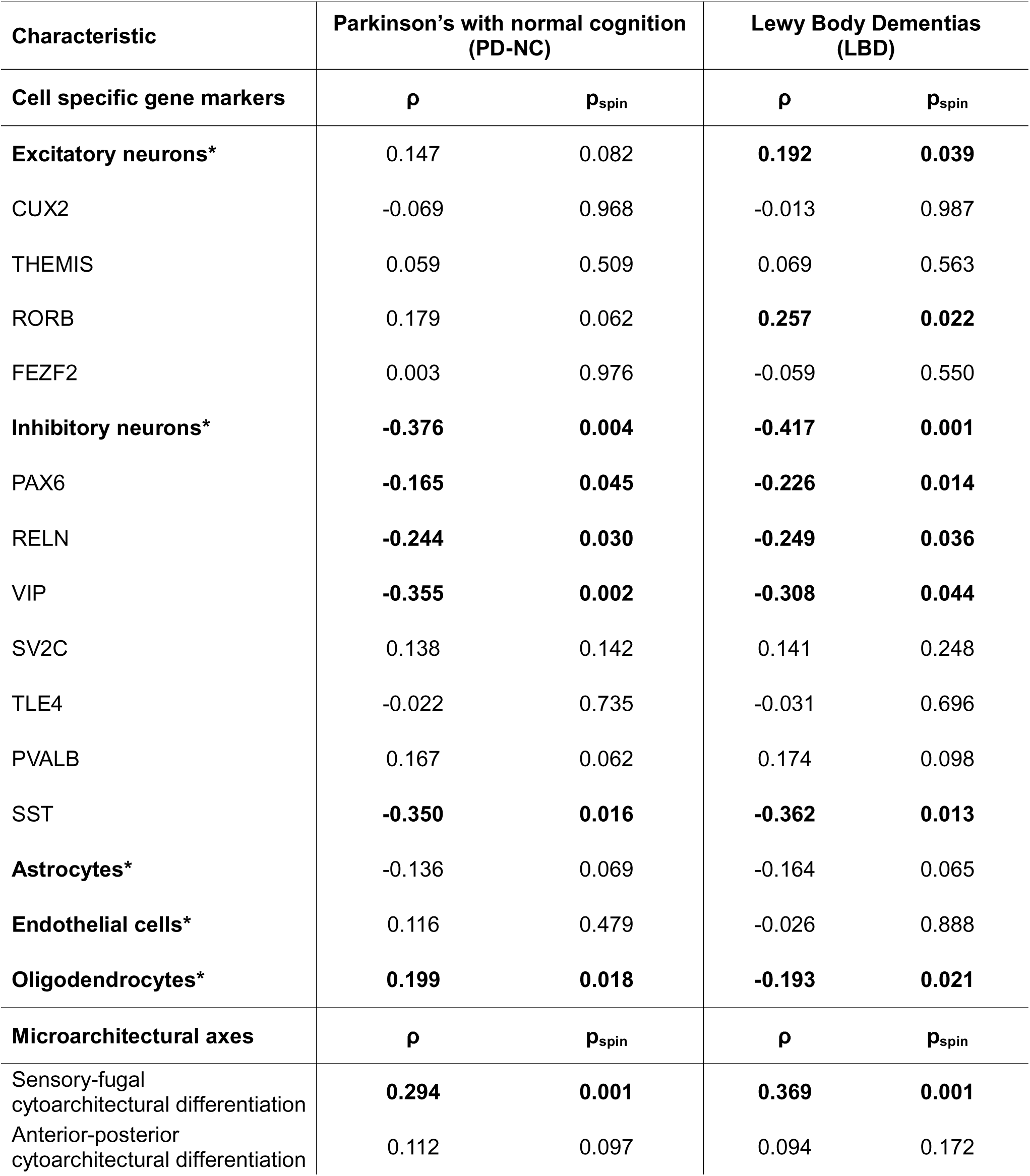
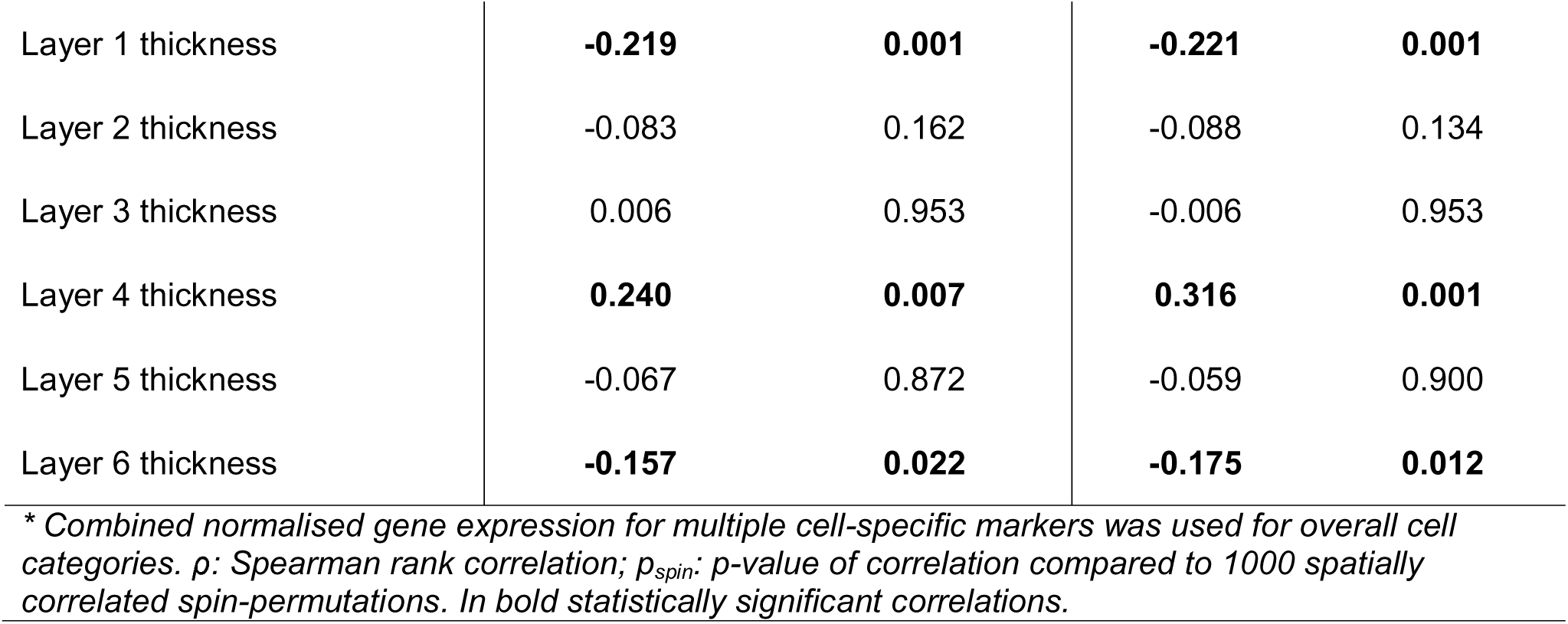
Neuronal and microstructural underpinnings of structural gradient 1 (SC-G1) alterations in Lewy Body Diseases.

| Characteristic | Parkinson's with normal cognition (PD-NC) |  | Lewy Body Dementias (LBD) |  |
| --- | --- | --- | --- | --- |
| Cell specific gene markers | $\rho$ | $p_{spin}$ | $\rho$ | $p_{spin}$ |
| **Excitatory neurons\*** | 0.147 | 0.082 | **0.192** | **0.039** |
| CUX2 | -0.069 | 0.968 | -0.013 | 0.987 |
| THEMIS | 0.059 | 0.509 | 0.069 | 0.563 |
| RORB | 0.179 | 0.062 | **0.257** | **0.022** |
| FEZF2 | 0.003 | 0.976 | -0.059 | 0.550 |
| **Inhibitory neurons\*** | **-0.376** | **0.004** | **-0.417** | **0.001** |
| PAX6 | **-0.165** | **0.045** | **-0.226** | **0.014** |
| RELN | **-0.244** | **0.030** | **-0.249** | **0.036** |
| VIP | **-0.355** | **0.002** | **-0.308** | **0.044** |
| SV2C | 0.138 | 0.142 | 0.141 | 0.248 |
| TLE4 | -0.022 | 0.735 | -0.031 | 0.696 |
| PVALB | 0.167 | 0.062 | 0.174 | 0.098 |
| SST | **-0.350** | **0.016** | **-0.362** | **0.013** |
| **Astrocytes\*** | -0.136 | 0.069 | -0.164 | 0.065 |
| **Endothelial cells\*** | 0.116 | 0.479 | -0.026 | 0.888 |
| **Oligodendrocytes\*** | **0.199** | **0.018** | **-0.193** | **0.021** |
| Microarchitectural axes | $\rho$ | $p_{spin}$ | $\rho$ | $p_{spin}$ |
| Sensory-fugal cytoarchitectural differentiation | **0.294** | **0.001** | **0.369** | **0.001** |
| Anterior-posterior cytoarchitectural differentiation | 0.112 | 0.097 | 0.094 | 0.172 |
| Layer 1 thickness | **-0.219** | **0.001** | **-0.221** | **0.001** |
| Layer 2 thickness | -0.083 | 0.162 | -0.088 | 0.134 |
| Layer 3 thickness | 0.006 | 0.953 | -0.006 | 0.953 |
| Layer 4 thickness | **0.240** | **0.007** | **0.316** | **0.001** |
| Layer 5 thickness | -0.067 | 0.872 | -0.059 | 0.900 |
| Layer 6 thickness | **-0.157** | **0.022** | **-0.175** | **0.012** |

\* Combined normalised gene expression for multiple cell-specific markers was used for overall cell categories. $\rho$ : Spearman rank correlation; $p_{spin}$ : p-value of correlation compared to 1000 spatially correlated spin-permutations. In bold statistically significant correlations.

### Microarchitectural underpinnings of structural gradient alterations

Next, we examined whether disease-related differences in the principal structural connectivity gradient reflect large-scale differences in cortical microstructure. We correlated changes in SC-G1 values (t-maps) in LBD vs controls and PD-NC vs controls to atlases of normative cytoarchitectural organisation and laminar thickness.

We found that primary gradient (SC-G1) alterations were spatially correlated with sensory-fugal cytoarchitectural differentiation derived from normative histology data, or a gradient of differentiation from more unimodal sensory areas to more transmodal cortical areas (LBD vs HC gradient changes: ρ=-0.369, p_spin_<0.001; PD-NC vs HC: ρ=0.295, p_spin_<0.001). This was replicated by using a unimodal to transmodal ranking based on the functional network allocation for each region (LBD: ρ=-0.352, p_spin_=0.001; PD-NC: ρ=-0.352, p_spin_=0.001) with more transmodal regions more likely to have increased gradient scores and more unimodal regions likely to have contracted gradient scores. In contrast there was no correlation with an anterior to posterior axis of cytoarchitectural differentiation (*Figure 3B*).

Differences in structural connectivity organisation in LBD and PD-NC also showed specific laminar contributions, with both SC-G1 gradient change t-maps showing negative correlations with regional distribution of layer 1 and 6 thickness and positive correlation with layer 4 thickness (*Table 2*). This suggests that regions with more reduced SC-G1 values in those groups are more likely to have thicker layer 1 and 6 and thinner layer 4.

### Structural gradient alterations are disease-specific and not driven by Alzheimer’s pathology

To assess whether the differences in structural cortical organisation we saw in LBD participants were driven by Alzheimer’s co-pathology or were disease-specific we performed two analyses. First, we assessed whether composite gradient difference scores were correlated to plasma p-tau217 levels (a plasma biomarker that corelates to brain levels of beta-amyloid and tau on PET^30^. We found no correlation for SC-G1 (ρ=0.132.09, p=0.346) or SC-G2 (ρ=0.172, p=0.217).

Secondly, we examined whether LBD-related differences in SC-G1 (unthresholded t-map of SC-G1 changes between LBD vs HC) were correlated with regional expression of mendelian risk genes for PD and AD, using spatially-correlated spin-permutations. We found that LBD gradient changes were correlated with expression of PD specific genes (ρ=-0.256, p_spin_=0.039) but not AD genes (ρ=-0.046, p_spin_=0.549). Together these findings suggest that Alzheimer’s co-pathology does not drive the gradient alterations seen in LBD participants.

### Structural gradient differences are associated with variation in regional gene expression patterns: the same regions drive this in LBD and PD-NC but there are differences in biological processes and pathways

Finally, we wanted to assess whether principal structural gradient alterations in LBD and PD-NC are underpinned by normative differences in genes related to specific biological processes and pathways. To do this, we performed partial least squares regression (PLS) with dependent variable Y the t-map of SC-G1 gradient alterations (1*200 regions, LBD vs HC and PD-NC vs HC) and predictor matrix X regional gene expression (17545*200 regions from the Allen Human Brain Atlas^31^); we tested our results against 10000 permutations of spatially correlated spin-permutations of the gradient t-map. The first principal component PLS1 explained the most variance in both gene expression and gradient alteration variability in both cases (*Supplementary Table 4*) and therefore PLS1 gene weightings were used for further gene ontology and enrichment analyses (only for genes significantly differentially weighted p_spin_<0.05).

For both LBD and PD-NC compared to controls, we observed that shifts in cortical gradients were associated with down-weighting, or reduced expression of specific genes: 2782 genes in LBD and 2296 genes in PD-NC. PLS1 region weights were positively correlated with gradient alterations seen in LBD (ρ=0.389, p_spin_<0.001). In other words, areas where gradient scores differed most in LBD had lower expression of genes with the most negative PLS1 weights. The regional profile of PLS1 weighting for both LBD and PD-NC was similar, with differences in regional expression of the same left parietal and frontal regions in both patient cohorts (*Figure 4A*).

**Figure 4.**
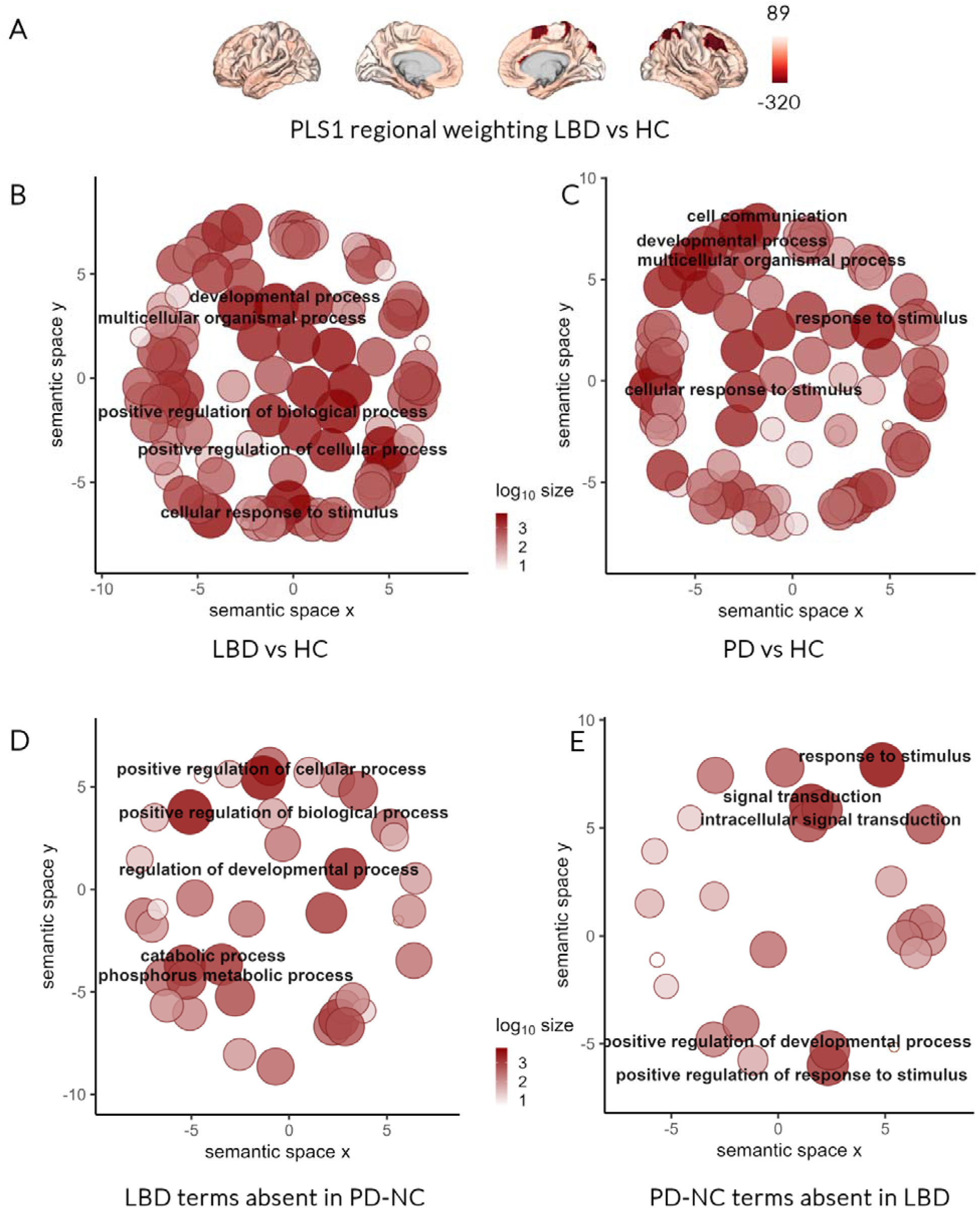
Structural gradient alterations in Lewy body disease are associated with regional gene expression patterns. We assessed whether principal structural gradient alterations in patients with Lewy body dementia (LBD) and Parkinson’s with intact cognition (PD-NC) are underpinned by normative differences in genes related to specific biological processes and pathways. We performed partial least squares regression (PLS) with dependent variable Y the t-map of primary structural gradient (SC-G1) alterations (1*200 regions, LBD vs HC and PD vs HC assessed separately) and predictor matrix X of regional gene expression (17545*200 regions derived from the Allen Human brain atlas^31^). The first principal component (PLS1), explaining most variance in both gene expression and SC-G1 variability was used in further analyses. The regional profile of PLS1 weighting for LBD vs GC **(A)** reveals that left parietal and frontal regions showed most downweighting (reduced expression) of genes. Genes that were significantly downweighted (against 1000 spatially correlated spin-permutations, p_spin_<0.05) were then included in gene ontology (GO) and enrichment analyses. Down-weighted genes, that were less expressed in regions showing SC-G1 differences in LBD vs HC, were enriched for GO terms relating to cellular response to stimulus, positive regulation of biological and cellular processes, and developmental processes **(B)** with similar results for down-weighted genes in PD-NC vs HC **(C)**. Although GO terms for LBD and PD-NC were highly inter-correlated, there were several terms that were solely enriched in LBD and not in PD-NC **(D)** or only enriched in PD-NC and not in LBD patients **(E)**.

Down-weighted genes, that were less expressed in regions showing differences in LBD vs controls, were enriched for GO terms relating to cellular response to stimulus, positive regulation of biological and cellular processes, and developmental processes (*Figure 4B*). They were also enriched for REACTOME pathways relating to lipid-associated metabolic pathways (*Table 3*).

**Table 3.** Most significantly enriched GO biological terms and REACTOME Pathways.

| GO Biological Terms |  |  |  |  |  |  |  |
| --- | --- | --- | --- | --- | --- | --- | --- |
| Parkinson's with normal cognition (PD-NC) |  |  |  | Lewy Body Dementias (LBD) |  |  |  |
| Term ID | Term Name | Term size | P-value | Term ID | Term Name | Term size | P-value |
| GO:0007267 | Cell-cell signalling | 1316 | 8.17E-12 | GO:0044281 | Small molecule metabolic process | 1795 | 1.34E-17 |
| GO:0044281 | Small molecule metabolic process | 1795 | 9.39E-12 | GO:0019752 | Carboxylic acid metabolic process | 905 | 6.02E-12 |
| GO:0006810 | Transport | 4341 | 1.30E-10 | GO:0006810 | Transport | 4341 | 1.26E-11 |
| GO:0006631 | Fatty acid metabolic process | 398 | 3.12E-10 | GO:0043436 | Oxoacid metabolic process | 927 | 1.41E-11 |
| GO:0065008 | Regulation of biological quality | 2847 | 1.19E-09 | GO:0006082 | Organic acid metabolic process | 933 | 2.26E-11 |
| GO:0051049 | Regulation of transport | 1589 | 3.82E-09 | GO:0006631 | Fatty acid metabolic process | 398 | 7.75E-11 |
| GO:0010646 | Regulation of cell communication | 3453 | 4.01E-09 | GO:0051049 | Regulation of transport | 1589 | 5.43E-10 |
| GO:0019752 | Carboxylic acid metabolic process | 905 | 7.27E-09 | GO:0065008 | Regulation of biological quality | 2847 | 6.65E-10 |
| GO:0032879 | Regulation of localization | 2005 | 8.08E-09 | GO:0007267 | Cell-cell signalling | 1316 | 1.67E-09 |
| GO:0023051 | Regulation of signalling | 3446 | 1.01E-08 | GO:0032879 | Regulation of localization | 2005 | 6.79E-09 |

| REAC Pathways |  |  |  |  |  |  |  |
| --- | --- | --- | --- | --- | --- | --- | --- |
| Parkinson's with normal cognition (PD-NC) |  |  |  | Lewy Body Dementias (LBD) |  |  |  |
| Term ID | Term Name | Term size | P-value | Term ID | Term Name | Term size | P-value |
| REAC:R-HSA-8978868 | Fatty acid metabolism | 173 | 7.50E-07 | REAC:R-HSA-8978868 | Fatty acid metabolism | 173 | 3.87E-06 |
| REAC:R-HSA-5661231 | Metallothioneins bind metals | 11 | 3.51E-04 | REAC:R-HSA-77289 | Mitochondrial Fatty Acid Beta-Oxidation | 37 | 8.45E-04 |
| REAC:R-HSA-77289 | Mitochondrial Fatty Acid Beta-Oxidation | 37 | 4.95E-04 | REAC:R-HSA-389887 | Beta-oxidation of pristanoyl-CoA | 8 | 9.19E-04 |
| REAC:R-HSA-389887 | Beta-oxidation of pristanoyl-CoA | 8 | 6.56E-04 | REAC:R-HSA-5661231 | Metallothioneins bind metals | 11 | 1.56E-03 |
| REAC:R-HSA-9033241 | Peroxisomal protein import | 60 | 8.84E-03 | **REAC:R-HSA-193368** | **Synthesis of bile acids and bile salts via 7alpha-hydroxycholesterol** | **23** | **4.44E-03** |
| REAC:R-HSA-5660526 | Response to metal ions | 14 | 9.17E-03 | REAC:R-HSA-8980692 | RHOA GTPase cycle | 149 | 1.35E-02 |
| REAC:R-HSA-390696 | Adrenoceptors | 9 | 1.26E-02 | **REAC:R-HSA-1430728** | **Metabolism** | **2079** | **1.36E-02** |
| **REAC:R-HSA-556833** | **Metabolism of lipids** | **735** | **2.14E-02** | REAC:R-HSA-9033241 | Peroxisomal protein import | 60 | 1.37E-02 |
| REAC:R-HSA-390918 | Peroxisomal lipid metabolism | 24 | 2.32E-02 | REAC:R-HSA-390696 | Adrenoceptors | 9 | 1.80E-02 |
| REAC:R-HSA-8980692 | RHOA GTPase cycle | 149 | 4.61E-02 | **REAC:R-HSA-164938** | **Nef-mediates down modulation of cell surface receptors by recruiting them to clathrin adapters** | **21** | **2.16E-02** |

*The GO and REAC term ID is a unique gene ontology identifier; the term name gives a brief description; and the term size is the number of unique genes associated with a given term. The reported P-values are corrected for multiple comparisons using the g:SCS algorithm in g:Profiler. In bold terms that are missing in the other respective patient group.*

We found a similar pattern for Gene Ontology (GO) terms and REACTOME pathways linked to down-weighted genes in PD-NC compared to controls (see Figure=4C). The ranking of GO terms for LBD vs HC and PD-NC vs HC was highly inter-correlated (ρ=0.807, p<0.001) with greater overlap than expected by chance (hypergeometric test p<0.001). The same was true for REACTOME pathways (ρ=0.733, p=0.025, hypergeometric test p<0.001). However, there were several terms that were exclusively enriched in one disease group (*Figure 4D-E*). 33 terms were only enriched in the PD-NC downweighted gene set and primarily involved signal transduction and response to stimulus. Whilst 62 terms were uniquely enriched in the LBD gene set and related to positive regulation of cellular processes, developmental processes and catabolic processes. Full tables of GO terms are seen in *Supplementary Table 5*.

## Discussion

Here, we comprehensively assessed cortical organisation differences in Lewy Body Diseases and how these changes relate to underlying cellular and molecular processes. We showed that differences in PD-NC and LBD diverged. In PD-NC we found expansion of cortical gradients, suggesting an exacerbation of interregional differences. In contrast, in LBD we found attenuation of interregional differentiation with overall gradient distribution consistent with controls; however, this was accompanied by widespread differences in cortical organisation of regional cortical gradient values in LBD patients. We show that these changes are relevant to disease severity and specific to cognitive but not motor severity.

Importantly, we were able to relate these differences in cortical hierarchies to normative cell type composition, cortical cytoarchitecture and gene expression. We found that cortical re-organisation in both PD-NC and LBD patients is more pronounced in regions that are normally richer in inhibitory neurons and poorer in oligodendrocytes. However, only differences in LBD patients were correlated with excitatory neurons, specifically RORB-expressing layer 4 neurons. Similarly, despite similarities in regional distribution of organisational alterations and in biological processes and pathways driving these between PD-NC and LBD participants, we found crucial differences. Cortical organisational differences in LBD but not PD-NC was more commonly seen in regions with reduced expression of genes related to signal transduction and response to stimuli; in PD-NC differences were more frequent in regions with reduced expression of genes related to positive regulation of biological and cellular processes and catabolism.

Together, our findings demonstrate 1) the association between disease-related gradient differences and cortical cytoarchitecture, neuronal populations and gene expression, 2) important differences between PD-NC and LBD suggesting different processes may underly cognitive and motor symptoms in Lewy body diseases and 3) the importance of multi-scale approaches integrating macroscale organisational principles with microarchitectural features for both validation and new hypothesis generation.

We found differences in cortical organisation in both PD-NC and LBD participants within the same brain regions, albeit with much more extensive differences at the regional level seen in LBD, but these had different overall effects on global cortical hierarchies. In PD-NC inter-regional differences appeared to be amplified, whereas in LBD overall gradient distribution was constricted, suggesting attenuation of regional specificity. Our findings replicate, in living patients, observations about regional cortical specialisation from post-mortem PD and PDD brains^29^. We now demonstrate this finding in-vivo and expand these observations to examine the whole-cortex organisational differences occurring in PD-NC and LBD. Importantly, we replicated the observation of divergent regional specialisation between PD-NC and LBD using two different cohorts, imaging modalities and analysis techniques.

Additionally, we replicated the observation found using single-cell transcriptomics of selective vulnerability of excitatory neurons in Lewy body dementias^29,32^, particularly RORB-expressing excitatory neurons^29^. In LBD patients, cortical gradient differences were positively correlated with normative regional expression of RORB, a gene specifically expressed in layer 4 excitatory neurons^33^, unlike in PD-NC. As gradient differences in LBD mostly reflected reduced gradient scores compared to controls, this implies that regions normally less rich in RORB (*Figure 2A)* and with thinner layer 4 (*Figure 2C*) are more likely to change within the cortical hierarchy in LBD. Layer 4, the internal granular layer, varies considerably in thickness and composition across different cortical areas and its small granular neurons are major postsynaptic targets of thalamic sensory nuclei and play a key role in sensory processing^33,34^. Altered sensory processing, specifically for visual perception, is a cornerstone of Lewy body diseases, with changes in visual processing happening early in the disease course or even predating symptoms^35,36^ and correlating with cognitive impairment, predicting the development of dementia in PD patients^5,37^. Higher divergence from the cortical organisation seen in controls was strongly correlated with cognitive but not motor scores and only in LBD participants but not in PD with normal cognition, suggesting these differences are specific to cognitive rather than motor impairment. A selective vulnerability of layer-4 neurons to pathological processes happening only in those patients with Lewy body disease with impaired cognition, could potentially explain both the visuoperceptual deficits seen in Lewy body disease, and why these deficits herald cognitive impairment^5,37^.

There is some evidence that vulnerability of layer 4 may be specific to alpha-synuclein pathology. Lewy bodies are more concentrated in lower cortical layers III to VI^38^ and although a study comparing post-mortem visual cortex between AD and DLB did not show major laminar specific differences, it did reveal functional changes within layer 4 GABAergic neurons^39^. Tau-pathology seems to spare layer 4 in Alzheimer’s^40^, with RORB-expressing layer 4 neurons particularly resilient in the presence of amyloid and tau pathology^41^. In contrast, gene expression changes within RORB-expressing excitatory neurons in PDD are explained by Lewy body but not amyloid pathological load^29^. Our findings that the regional distribution of gradient alterations in LBD was correlated with the regional distribution of PD but not AD specific gene lists and that divergence from normative gradient ratings did not correlate with plasma p-tau levels provide further evidence that the differences in cortical organisation we show in LBD patients may not only be specific to cognition but also to alpha-synuclein pathology.

Whilst RORB layer 4 excitatory neurons may play a differential role in dementia in LBD, we also found important similarities between the drivers of gradient differences in LBD and PD-NC. In both patient groups, differences in cortical organisation followed the sensori-fugal axis, with primary sensory areas more downweighted; and this correlated with inhibitory neuron and oligodendrocyte regional expression. Impaired intracortical inhibition is well described in relation to motor symptoms in PD^42–44^. And while PD was traditionally considered as a purely neuronal disease, there is increasing evidence that oligodendrocytes play a significant role in the early stages of the disease: transcriptomic changes in oligodendrocytes are repeatedly described in cortical and deep regions post-mortem^45–47^ and structural changes within the white matter, before or in the absence of grey matter loss is seen repeatedly in animal and cell models^48–50^ as well as using neuroimaging in in-vivo brains^11,12,51^. Our findings provide further evidence for this, albeit not supporting a specificity of oligodendrocyte involvement for cognitive rather than motor impairment.

Despite the striking structural organisational differences we saw in PD-NC and LBD compared to controls, we found no changes in functional gradients. Although this could reflect reduced power from a smaller sample size due to rsfMRI scans not passing quality control, this could reflect either more localised changes or compensatory mechanisms.

Functional connectivity compensation may play a role in delaying symptom onset during the presymptomatic phase and its loss may partially explain motor and cognitive symptom severity in Parkinson’s^52–54^. Although we were not able to assess compensation in this study, this could be clarified in future work.

The concordance of our in-vivo results with transcriptional alterations seen in PD and PDD at post-mortem^29^ suggests that, at least partly, differences in cortical hierarchies that we identified using gradient analyses are driven by changes in gene expression in Lewy body diseases. It also provides important support for the use of multimodal and multiscale approaches combining neuroimaging with atlases of normative gene expression and cytoarchitecture. Although the value of such approaches in bridging scales and providing mechanistic insights into the healthy human brain is well-established^13,14,16^, ongoing debate on their value and applicability in the presence of disease persists^55,56^, not only due to methodological limitations but also because gene expression and cell composition is altered in neurodegeneration^29^. However, our findings suggest that these methods can yield important insights and can be used in a bidirectional way to inform experiments of gene expression: not only can these methods be used to validate experiments at the cellular or molecular level as we have done, but they can also generate new hypotheses and calibrate these experiments. For example, future work aimed to assess transcriptomic changes and how these differ between LBD and PD could focus on sampling those regions where we found that the contribution of gene expression variability on cortical re-organisation to be highest (PLS1 weightings, *Figure 3A*); whilst layer 4 specific differential changes in LBD versus Alzheimer’s disease could focus on sensory areas and particularly the sensorimotor cortex which was also an area significantly downweighted in LBD (and thus correlated with reduced RORB expression and layer 4 thickness).

In summary, our study suggests that changes at the molecular level within the same regions lead to divergent alterations in cortical organisation in patients with Lewy body diseases with and without dementia. Despite similarities in the cellular and molecular drivers of organisational differences between those with and those without cognitive impairment, we also show important differences particularly in RORB layer 4 excitatory neurons; these are not driven by amyloid co-pathology and are specific to cognition. Finally, our study provides an important example of how multi-scale in-vivo neuroimaging can be used to provide fundamental insights into the molecular underpinnings of neurodegeneration.

## Acknowledgements

AZ is supported by an Alzheimer’s Research UK Clinical Research Fellowship (2018B-001) and grants from Parkinson’s UK and Rosetrees Trust/Race Against Dementia. RSW is supported by a Wellcome Career Development Award (205263/Z/22/Z). MGP and MR are funded by Aligning Science Across Parkinson’s (ASAP-000478 and ASAP-000509) through the Michael J. Fox Foundation for Parkinson’s Research (MJFF). This work was also supported by grants from the Lewy Body Society, Rosetrees Trust, an Academy of Medical Sciences Starter Grant, the National Institute for Health Research University College London Hospitals Biomedical Research Centre and the UK Dementia Research Institute UKDRI-2206 through UK DRI Ltd, principally funded by the Medical Research Council. RSW has received honoraria from GE Healthcare, Bial, Omnix Pharma and Britannia and consultancy fees from Therakind and Accenture, and UCL Partners.

## Code availability

All analysis code will be made available upon publication: https://github.com/AngelikaZa/RegionalDifferentiationLBD.

## Methods

### Participants

For both 3T and 7T cohorts, participants were recruited to University College London (UCL). All participants provided written informed consent, and the study was approved by the Queen Square Research Ethics Committee (15.LO.0476). The 3T cohort has been previously described^5,11^. In brief, participants were required to be over 50 years of age, capable of providing informed consent and able to comply with study procedures. Additional inclusion criteria included:

- *Participants with Parkinson’s disease (PD):* a clinical diagnosis in accordance with the Movement Disorders Society (MDS) clinical diagnostic criteria^57^.
- *Participants with Dementia with Lewy Bodies (DLB)*: a clinical diagnosis of probable DLB in accordance with the McKeith diagnostic criteria^2^.
- *Participants with Parkinson’s disease dementia (PDD):* an established clinical diagnosis of Parkinson’s disease and dementia or impaired function in activities of daily leaving (impaired functional assessments questionnaire) and a Montreal Cognitive Assessment (MoCA) score below 26^3^.
- *Participants with Parkinson’s disease and mild cognitive impairment (PD-MCI):* a clinical diagnosis of PD^57^ and persistent performance below 1.5 standard deviations (SD) in at least two different tests in one cognitive domain or one cognitive test in at least two cognitive domains, according to respective MDS clinical criteria^58^.
- *Control participants:* aged between 50 and 81.

Exclusion criteria included confounding neurological and psychiatric disorders and contraindications to MRI. Controls diagnosed with dementia, or mild cognitive impairment or Mini-mental State Examination (MMSE) score less than 25 were also excluded.

Participants with a diagnosis of DLB, PDD or PD-MCI were grouped as Lewy Body Dementia (LBD), with the remaining PD participants classified as PD-normal cognition (PD-NC).

#### Replication cohort

Due to stricter rules for inclusion for 7T MRI safety, we did not restrict participants with PD or LBD in terms of disease duration. Otherwise, diagnostic categories and inclusion-exclusion criteria were identical to that of the main 3T cohort.

A total of 46 (n=21 controls, n=13 PD-NC and 13 LBD) participants with scans passing quality assurance (see *MRI quality control* below) were included. Of those participants, n=23 (50%) also had 3T scanning and are included in the 3T cohort. For those participants that had both a 3T and 7T scan, scanning for 7T was performed between 1 and 4 years after their 3T scan.

#### Clinical assessments

All participants underwent the same clinical and neuropsychological assessments. All assessments were performed with participants receiving their usual medications to minimize discomfort and minimise attrition. The Mini-Mental State Examination^59^ (MMSE) and Montreal Cognitive Assessment^60^ (MoCA) were used as measures of global cognition and 2 tests per cognitive domain were performed^58^:

- *Attention*: Stroop colour naming task^61^ and Digit span backwards^62^
- *Executive function*: Category fluency^63^ and Stroop interference task^61^
- *Language*: Letter fluency^63^ and Graded naming task^64^
- *Memory*: Word recognition task^65^ and Logical memory task (delayed score) ^62^
- *Visuospatial function*: Hooper visual organization test^66^ and Benton Judgement of line orientation^67^.

A composite cognitive score was calculated as the averaged z-scores of the MoCA plus one task per cognitive domain: inverted Stroop (colour naming time), Category fluency, Letter fluency, Word recognition task and Hooper Visual Organization Test, as we have previously described^5,11^.

Global disease burden was assessed using the MDS United Parkinson’s Disease Rating Scale (UPDRS) total score^68^. Motor severity was assessed using the UPDRS part 3 (UPDRS-III)^68^, and the timed up and go test (TUG)^69^. Cognitive fluctuations were measured using the clinician assessment of fluctuations (CAF)^70^, one-day fluctuations scale^70^ and the dementia cognitive fluctuation scale (DCFS)^71^. Anxiety and depression were measured using the Hospital Anxiety and Depression Scale (HADS)^72^. Impairments in activities of daily living were measured using the Functional Activities Questionnaire^73^. Autonomic symptoms were assessed using the Compass-31 questionnaire^74^. Sleep disturbances were measured using the Rapid Eye Movement Behaviour Disorder Sleep Questionnaire (RBDSQ)^75^ and visual hallucinations using the University of Miami PD hallucinations questionnaire (UMPDHQ) ^76^. Levodopa equivalent daily doses (LEDD) were calculated for all PD-NC and LBD participants^77^.

#### Statistical analysis

Between group-comparisons for demographics and clinical characteristics were performed using ANOVA for normally distributed and Kruskal-Wallis tests for non-normally distributed variables, with post-hoc t-test and Mann-Whitney respectively. Shapiro-Wilk was used to determine normality.

### MRI data acquisition and quality control

All participants were scanned in the same scanners (3T Siemens Prisma scanner for main cohort and 7T Siemens Terra scanner for replication cohort) with the same protocol.

Neuroimaging included:

**Main cohort (3T):** structural T1-weighted scan (MPRAGE), diffusion-weighted imaging (DWI) and resting state functional MRI (rsfMRI).

DWI was acquired with the following parameters: b0 (both AP and PA directions), *b*=50 s/mm^2^/17 directions, *b*=300 s/mm^2^/8 directions, *b*=1000 s/mm^2^/64 directions, b=2000 s/mm^2^/64 directions, 2×2×2 mm isotropic voxels, TR=3260 ms, TE=58 ms, 72 slices, acceleration factor =2, acquisition time ∼10 min. 3D MPRAGE (magnetization prepared rapid acquisition gradient echo) image was acquired with: 1×1×1 mm isotropic voxels, TE=3.34 ms, TR=2530 ms, flip angle=7°, acquisition time ∼6min. rsfMRI was acquired with: gradient-echo EPI, TR=3.36 s, TE=30 ms, flip angle=90°, FOV=192×192, voxel size = 3×3×2.5 mm, 105 volumes, acquisition time ∼6 min. During rsfMRI, participants were instructed to lie quietly with eyes open and avoid falling asleep (confirmed by monitoring and post-scan debriefing).

**Replication cohort (7T):** structural T1-weighted scan (MP2RAGE) and multiparameter mapping (MPM) acquisitions.

All 7T data were acquired on the same 7T Siemens Terra Scanner with a multi-echo variable flip angle MPM protocol^78,79^ with whole-brain coverage and 0.6 mm isotropic voxel size. Two multi-echo 3D fast low angle shot (FLASH) scans with proton density and T1-weighting were acquired with flip angles 6° and 24° respectively, and readouts of alternating polarity to give 6 equidistant echoes with TE=2.2-14.1ms for PD and 2.3-14.2ms for T1 and a common TR of 19.5ms, FOV=256x218x173mm^3^, and readout bandwidth was 469 Hz/pixel. Parallel imaging was used with an acceleration factor of 2 in each phase and partition directions and 48 integrated reference lines. An additional multi-echo scan with magnetisation transfer weighting was acquired using the same settings as the PD weighted acquisition, but with just four echoes (2.2-9.34ms). This scan included a 180^0^ pre-pulse with 4ms duration, 2kHz off resonance. Calibration data to correct for transmit and receive field inhomogeneities were also acquired^80,81^.

Parameter maps were then calculated used the hMRI toolbox in SPM12 using default toolbox configuration settings with correction for imperfect spoiling enabled^79^ and the pre-computed B1 option. The following parameter maps were derived: proton density, sensitive to tissue water content, longitudinal relaxation rate (R1) sensitive to myelin, iron and water content, effective transverse relaxation rate (R2*) sensitive to iron content and less so myelin, and magnetisation transfer saturation (MTsat) particularly sensitive to myelin content.

All imaging sequences were performed with PD and LBD participants receiving their normal medications.

#### MRI quality control

Raw image data were visually inspected, by two raters blinded to clinical data, to ensure appropriate brain coverage and identify artefacts, e.g. motion. Further visual inspection was performed throughout the processing, including the final maps computed. In addition to visual inspection, we adopted strict motion-control criteria for rsfMRI, given susceptibility to motion artefact, similar to our previous work^28^. Only scans passing quality control for each modality were included. This resulted in a total of 111 participants included in the 3T functional connectivity analyses (n=23 controls, n=35 PD-NC, and n=50 LBD). Full details on MRI quality control and preprocessing of 3T data is seen in *Supplementary Methods*.

### Cortical gradient construction

#### Gradient construction

Following standard preprocessing of T1-weighted, DWI and rsfMRI data, using established pipelines in our group (see *Supplementary Methods*)^28^, structural and functional connectomes were constructed for each participant using the same parcellation (Shaefer 200 cortical regions)^82^. Parcellations in the range of 200 nodes increase reliability in gradient construction^83^, particularly those derived from functional connectivity. The same parcellation was used to construct both structural and functional connectomes, weighted by streamline count for structural and correlation coefficient for functional connectomes.

Cortical gradients were then derived separately from structural and functional connectivity matrices as well as from fused gradients using both structural and functional connectivity^14^. Gradient analyses were performed using BrainSpace^83^. Gradients were derived using diffusion map embedding which identifies spatial axes of variation in connectivity across different brain regions, whereby cortical vertices that are strongly interconnected are closer together and vertices with little or no inter-connectivity are farther apart^16,84^. Normalised angle was used as a metric of similarity for this at it retains angular distance between regions and is less sensitive to noise (values range between 0 and 1, with 1 denoting identical angles, and 0 opposing angles). The normalised angle between two nodes i and j (A(i,j)) is calculated as:

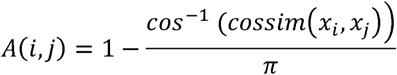

where *cossim* is the cosine similarity function and x is the respective connectivity metric (streamline count for structural and pearson correlation for functional connectivity matrices).

First, we generated a group-level gradient component template for each gradient type (structural, functional, fused) limiting the number of gradients to 10 and using default sparsity, thus keeping the top 10% of weights. Each group-level template was calculated from the average structural, functional and fused connectivity matrices of control participants (n=23). Then individual gradients were derived from each participant’s structural, functional and fused connectivity matrix (10 gradients, default sparsity 0.9, a=0.5 diffusion). Procrustes alignment was then applied to the gradient components of each individual to align them to the group template; this enables gradients to be compared across individuals or groups^85^. Results were replicated using different sparsity parameters (sparsity 0.8 and 0.5).

#### Statistical analysis

Gradient scores were compared between groups (PD-NC vs HC, LBC vs HC and LBD vs PD-NC) using surface-based linear models implements in BrainStat^86^, with age and sex as covariates and family-wise error (FWE) correction using random field theory and the default cluster-defining threshold 0.01.

### Neural contextualisation

We contextualised the differences in structural gradients found in LBD and PD-NC compared to HC with respect to normative variations in 1) excitatory and inhibitory neuronal gene expression markers, 2) cortical cytoarchitecture, 3) disease specific genetic and plasma markers, and 4) global gene expression.

#### Cell-specific gene expression

The unthresholded t-map of differences in structural gradients (LBD vs HC and PD vs HC), corrected for age and sex, was parcellated (Shaefer 200 cortical regions) using the neuromaps Parcellater tool^87^.

Gene expression profiles were obtained using data from the Allen Human Brain Atlas (AHBA)^31^. We extracted gene expression data and mapped them to 200 cortical regions using abagen and an established preprocessing pipeline^88,89^. Each tissue sample was assigned to an anatomical structure of the 200 cortical regions, using the AHBA MRI data for each donor. Data were pooled between homologous cortical regions to ensure adequate coverage of left (six donors with data) and right hemisphere (two donors with data).

Distances between samples were evaluated on the cortical surface with a 2 mm distance threshold. Probe-to-gene annotations were updated in Re-Annotator^90^ with a background threshold of 50% of samples. A representative probe for a gene was selected based on highest intensity. Gene expression data were normalised across the cortex using scaled, outlier-robust sigmoid normalisation. 15745 genes (of 20,737 initially included) survived preprocessing. The resulting gene table was used to extract cell-specific gene markers and in assessing global gene expression patterns.

Cell type specific markers included:

- *Excitatory neurons*: CUX2, THEMIS, RORB, FEZF2
- *Inhibitory neurons*: PAX6, RELN, VIP, SV2C, TLE4, PVALB, SST
- *Astrocytes*: AQP4, GFAP
- *Endothelial cells*: CLDN5, SEMA3G, EFNB2, MFSD2A, SLC16A1, C1QA, HBB, ACTA2, CNN1, VWF, TSHZ2
- *Microglia*: CD74, PTPRC
- *Oligodendrocytes*: PDGFRA, BCAS1, PLP1, MBP, RASGRF1, ANKRD18A

Pooled gene expression for each of the above cell types was calculated per parcel by adding normalised gene expression values for each individual gene. Regional cell-specific gene expression was compared to the parcellated gradient difference t-map using spearman correlation and spatial permutations (1000 spin permutations, statistical significance threshold p_spin_ <0.05).

#### Cytoarchitecture

Histology-derived gradient maps reflecting axes of differentiation on cytoarchitecture in the healthy human brain and average regional laminar thickness values were obtrained from BigBrain^91,92^; these are publicly available through BigBrainWarp^93^. Two main axes of differentiation were assessed: sensory-fugal (Histological gradient 1), differentiating between sensory and motor cortices from paralimbic structures and anterior-posterior (Histological gradient 2). We compared the parcellated t-map of gradient differences to these axes of cytoarchitectural differentiation, and to regional laminar thickness using spearman correlation and spin permutation tests (1000 permutations, p_spin_ <0.05).

#### Association with disease severity and specificity

To examine whether differences in cortical organisation are related to disease severity and specific for alpha-synuclein rather than driven by amyloid co-pathology we performed two analyses. First, we calculated for each participant, a composite gradient difference score. This was derived from the Z-scored value for SC-G1 for each region compared to the control SC-G1 value for that region; this was then summed across the 200 regions to a single composite gradient difference score *Gs*:

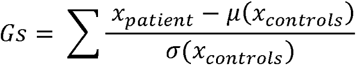

where x is the gradient value for a given region, μ is the mean and σ the standard deviation.

We correlated that score with outcomes of disease severity within each disease group (LBD and PD-NC) and with plasma p-tau217 using spearman correlation.

Secondly, we examined whether the spatial distribution of LBD-related SC-G1 differences (unthresholded t-map of SC-G1 differences between LBD vs HC) was correlated with the spatial pattern of regional expression for mendelian risk genes for PD and AD. We curated gene lists for Mendelian disease-associated genes from Blauwendraat et al^94^ for PD (including only those classified as high or very high confidence for PD causation) and from OpenTargets for Alzheimer’s disease, filtering for associated genes and requiring a genetic association score of >0.6. Our disease specific gene lists are in *Supplementary Methods*. We then extracted summed normalised gene expression values for each of the 200 cortical regions of the Schaeffer parcellation from the processed AHBA gene expression table above. We used spatially-correlated spin-permutations (1000 permutations, p_spin_ <0.05) to compare SC-G1 changes with regional disease-specific gene expression.

#### Regional patterns of gene expression

We used partial least squares regression (PLS) with predictor matrix X (the 200*15745 matrix of 200 regional mRNA measurements from 15475 genes extracted from AHBA^31^ as described above) and dependent variable Y the 200*1 regional structural gradient changes (t-map of gradient changes for LBD vs HC and PD vs HC separately). The first PLS component (PLS1) which explained the highest variance in gradient change and gene expression was used to weigh and rank gene predictor variables. 1000 spin permutations^95^ of the t-map of gradient changes was used to test the null hypothesis that PLS1 explained no more variance in *Y* than chance and the null hypothesis of zero weight for each gene (q<0.05 1000 permutations based on sphere-projection-rotations^95^). Genes with significantly different weights than expected by chances were included in subsequent enrichment analyses as in previous work^96–99^.

##### Enrichment analysis

Gene Ontology (GO), Kyoto Encyclopedia of Genes and Genomes (KEGG) and Reactome pathway databases were used as references for enrichment. Enrichment analysis was performed using g:Profiler^100^ with significance threshold p<0.05 (g:SCS method for multiple comparisons) and discarding terms associated with >2500 genes as being too general. The reduce and visualised gene ontology tool REViGO, based on semantic similarity, was used to visualise significant GO terms^101^. We used both rank spearman correlation and hypergeometric test to assess term similarity between LBD and PD-NC enriched terms.

## 7T#Region of interest analysis

MPM maps were generated using the hMRI toolbox^102^. Each participant’s MP2RAGE image was co-registered to the proton density image, and all subsequent processing was performed in each participants native proton density space. The same Schaffer parcellation (used for gradient construction in the 3T dataset) was used to parcellate the brain into 200 cortical regions. Four regions of interest were selected for analysis based on their SC-G1 rankings and whether they showed differences or not in their ranking between LBD vs HC. These comprised two regions from the extremes of the gradient ranking which showed differences in ranking between LBD and controls (“RH_SalVentAttn_TempOccPar_3”, and “RH_SomMot_18”) and two from the middle of the gradient distribution that did not show significant differences between LBD and HC (“RH_Default_Temp_1”, “LH_Default_Temp_1”). For each ROI we extracted mean MPM signal for each of the MPM maps (R1, R2*, proton density and MTsat) and constructed general linear models with these as the dependent variables. The models included main effects of ROI and Group (LBD, PD-NC, HC) as well as their interaction. Age and sex were additionally included as covariates of no interest. The effect of interest was the ROI*Group interaction, to assess if the MPM values in the ROIs depended on having LBD.

### Replication and robustness

Several replication analyses were performed to ensure robustness of results:

- *<u>Replication in 7T cohort using quantitative MRI</u>*: We used a separate cohort of ultra-high field 7T MRI to validate differences in inter-regional differentiation in LBD and PD-NC than controls. This used different imaging sequences, different patients and different analysis methods, further adding to the robustness of our results.
- *Gradient construction*: To ensure the choice of gradient construction did not influence results, we replicated our analyses with different sparsity options (sparsity 0.9 is presented in the main manuscript, and sparsity 0.8, and 0.5 are presented in *Supplementary Results*).
- *<u>Excitatory and inhibitory neuronal contribution</u>:* Two different analyses were used to assess the relationship between organisational differences seen in LBD (t-map of gradient change) and PD vs controls to different neuronal populations: 1) cell-specific markers for excitatory and inhibitory neurons and 2) PET density data for excitatory and inhibitory neurotransmitters (glumamate and GABA respectively). Therefore, we used different sources for normative variation in excitatory and inhibitory neuronal distribution derived from different populations and different modalities, increasing the robustness of our findings.
- *<u>Spatial autocorrelation:</u>* For all analyses of neural contextualisation we used spatial permutations, based on spatially correlated sphere rotations^95^, to test statistical significance. This ensures that correlations in the regional pattern of difference in cortical organisation (structural gradient differences seen in LBD vs controls and PD vs controls) and gene expression controls for false positive bias due to gene-gene co-expression and spatial autocorrelation^103^.
- *Enrichment analyses:* We only included significantly differentially weighted genes compared to spin permutations in enrichment analyses.

## References

1. Goedert, M., Spillantini, M. G., Del Tredici, K. & Braak, H. 100 years of Lewy pathology. Nat. Rev. Neurol. 9, 13–24 (2013).

2. McKeith, I. G. et al. Diagnosis and management of dementia with Lewy bodies. Neurology 89, 88–100 (2017).

3. Emre, M. et al. Clinical diagnostic criteria for dementia associated with Parkinson’s disease. Mov. Disord. Off. J. Mov. Disord. Soc. 22, 1689–1707; quiz 1837 (2007).

4. Williams-Gray, C. H. et al. The CamPaIGN study of Parkinson’s disease: 10-year outlook in an incident population-based cohort. J. Neurol. Neurosurg. Psychiatry 84, 1258–1264 (2013).

5. Hannaway, N. et al. Visual dysfunction is a better predictor than retinal thickness for dementia in Parkinson’s disease. J. Neurol. Neurosurg. Psychiatry (2023) doi:10.1136/jnnp-2023-331083.

6. Colloby, S. J., Watson, R., Blamire, A. M., O’Brien, J. T. & Taylor, J.-P. Cortical thinning in dementia with Lewy bodies and Parkinson disease dementia. Aust. N. Z. J. Psychiatry 54, 633–643 (2020).

7. Bhome, R. et al. A neuroimaging measure to capture heterogeneous patterns of atrophy in Parkinson’s disease and dementia with Lewy bodies. NeuroImage Clin. 42, 103596 (2024).

8. Mak, E. et al. Cortical microstructural abnormalities in dementia with Lewy bodies and their associations with Alzheimer’s disease copathologies. Npj Park. Dis. 11, 1–15 (2025).

9. Schumacher, J. et al. Free water imaging of the cholinergic system in dementia with Lewy bodies and Alzheimer’s disease. Alzheimers Dement. 19, 4549–4563 (2023).

10. Chiu, S. Y. et al. Longitudinal Free-Water Changes in Dementia with Lewy Bodies. Mov. Disord. Off. J. Mov. Disord. Soc. (2024) doi:10.1002/mds.29763.

11. Zarkali, A. et al. Neuroimaging and plasma evidence of early white matter loss in Parkinson’s disease with poor outcomes. Brain Commun. 6, fcae130 (2024).

12. Hannaway, N. et al. Neuroimaging and plasma biomarker differences and commonalities in Lewy body dementia subtypes. 2025.01.24.634687 Preprint at 10.1101/2025.01.24.634687 (2025).

13. Bernhardt, B. C., Smallwood, J., Keilholz, S. & Margulies, D. S. Gradients in brain organization. NeuroImage 251, 118987 (2022).

14. Paquola, C. et al. Microstructural and functional gradients are increasingly dissociated in transmodal cortices. PLOS Biol. 17, e3000284 (2019).

15. Burt, J. B. et al. Hierarchy of transcriptomic specialization across human cortex captured by structural neuroimaging topography. Nat. Neurosci. 21, 1251–1259 (2018).

16. Margulies, D. S. et al. Situating the default-mode network along a principal gradient of macroscale cortical organization. Proc. Natl. Acad. Sci. U. S. A. 113, 12574–12579 (2016).

17. Xia, Y. et al. Development of functional connectome gradients during childhood and adolescence. Sci. Bull. 67, 1049–1061 (2022).

18. Dong, H.-M., Margulies, D. S., Zuo, X.-N. & Holmes, A. J. Shifting gradients of macroscale cortical organization mark the transition from childhood to adolescence. Proc. Natl. Acad. Sci. 118, e2024448118 (2021).

19. Valk, S. L. et al. Genetic and phylogenetic uncoupling of structure and function in human transmodal cortex. Nat. Commun. 13, 2341 (2022).

20. Valk, S. L. et al. Shaping brain structure: Genetic and phylogenetic axes of macroscale organization of cortical thickness. Sci. Adv. 6, eabb3417 (2020).

21. Burt, J. B. et al. Hierarchy of transcriptomic specialization across human cortex captured by structural neuroimaging topography. Nat. Neurosci. 21, 1251–1259 (2018).

22. Fornito, A., Arnatkevičiūtė, A. & Fulcher, B. D. Bridging the Gap between Connectome and Transcriptome. Trends Cogn. Sci. 23, 34–50 (2019).

23. Xia, M. et al. Connectome gradient dysfunction in major depression and its association with gene expression profiles and treatment outcomes. Mol. Psychiatry 27, 1384–1393 (2022).

24. Holmes, A. et al. Disruptions of Hierarchical Cortical Organization in Early Psychosis and Schizophrenia. Biol. Psychiatry Cogn. Neurosci. Neuroimaging 8, 1240–1250 (2023).

25. Dong, D. et al. Compressed sensorimotor-to-transmodal hierarchical organization in schizophrenia. Psychol. Med. 53, 771–784 (2023).

26. Royer, J. et al. Cortical microstructural gradients capture memory network reorganization in temporal lobe epilepsy. Brain 146, 3923–3937 (2023).

27. Li, J. et al. Morphometric brain organization across the human lifespan reveals increased dispersion linked to cognitive performance. PLOS Biol. 22, e3002647 (2024).

28. Zarkali, A. et al. Organisational and neuromodulatory underpinnings of structural-functional connectivity decoupling in patients with Parkinson’s disease. *Commun*. Biol. 4, 1–13 (2021).

29. Fairbrother-Browne, A. et al. Molecular and cellular signatures differentiate Parkinson’s disease from Parkinson’s disease with dementia. 2025.03.04.641379 Preprint at 10.1101/2025.03.04.641379 (2025).

30. Kivisäkk, P. et al. Clinical evaluation of a novel plasma pTau217 electrochemiluminescence immunoassay in Alzheimer’s disease. Sci. Rep. 14, 629 (2024).

31. Hawrylycz, M. et al. Canonical genetic signatures of the adult human brain. Nat. Neurosci. 18, 1832–1844 (2015).

32. Goralski, T. M. et al. Spatial transcriptomics reveals molecular dysfunction associated with cortical Lewy pathology. Nat. Commun. 15, 2642 (2024).

33. Siletti, K. et al. Transcriptomic diversity of cell types across the adult human brain. Science 382, eadd7046 (2023).

34. Pluta, S. et al. A direct translaminar inhibitory circuit tunes cortical output. Nat. Neurosci. 18, 1631–1640 (2015).

35. Han, G. et al. Visual Acuity and Development of Parkinson’s Disease: A Nationwide Cohort Study. Mov. Disord. 35, 1532–1541 (2020).

36. Hamedani, A. G., Abraham, D. S., Maguire, M. G. & Willis, A. W. Visual Impairment Is More Common in Parkinson’s Disease and Is a Risk Factor for Poor Health Outcomes. Mov. Disord. Off. J. Mov. Disord. Soc. (2020) doi:10.1002/mds.28182.

37. Zarkali, A., McColgan, P., Leyland, L.-A., Lees, A. J. & Weil, R. S. Visual Dysfunction Predicts Cognitive Impairment and White Matter Degeneration in Parkinson’s Disease. Mov. Disord. n/a, (2021).

38. Harding, A. J. & Halliday, G. M. Simplified neuropathological diagnosis of dementia with Lewy bodies. Neuropathol. Appl. Neurobiol. 24, 195–201 (1998).

39. Khundakar, A. A. et al. Analysis of primary visual cortex in dementia with Lewy bodies indicates GABAergic involvement associated with recurrent complex visual hallucinations. Acta Neuropathol. Commun. 4, 66 (2016).

40. Duyckaerts, C., Delatour, B. & Potier, M.-C. Classification and basic pathology of Alzheimer disease. Acta Neuropathol. (Berl.) 118, 5–36 (2009).

41. Dharshini, S. A. P. et al. Molecular Signatures of Resilience to Alzheimer’s Disease in Neocortical Layer 4 Neurons. 2024.11.03.621787 Preprint at 10.1101/2024.11.03.621787 (2024).

42. Berardelli, A., Rona, S., Inghilleri, M. & Manfredi, M. Cortical inhibition in Parkinson’s disease: A study with paired magnetic stimulation. Brain 119, 71–77 (1996).

43. Ammann, C. et al. Cortical disinhibition in Parkinson’s disease. Brain 143, 3408– 3421 (2020).

44. Ridding, M. C., Rothwell, J. C. & Inzelberg, R. Changes in excitability of motor cortical circuitry in patients with parkinson’s disease. Ann. Neurol. 37, 181–188 (1995).

45. Bryois, J. et al. Genetic identification of cell types underlying brain complex traits yields insights into the etiology of Parkinson’s disease. Nat. Genet. 52, 482–493 (2020).

46. Agarwal, D. et al. A single-cell atlas of the human substantia nigra reveals cell-specific pathways associated with neurological disorders. Nat. Commun. 11, 4183 (2020).

47. Smajić, S. et al. Single-cell sequencing of human midbrain reveals glial activation and a Parkinson-specific neuronal state. Brain J. Neurol. 145, 964–978 (2022).

48. Volpicelli-Daley, L. A. et al. Exogenous α-Synuclein Fibrils Induce Lewy Body Pathology Leading to Synaptic Dysfunction and Neuron Death. Neuron 72, 57–71 (2011).

49. Chung, C. Y., Koprich, J. B., Siddiqi, H. & Isacson, O. Dynamic changes in presynaptic and axonal transport proteins combined with striatal neuroinflammation precede dopaminergic neuronal loss in a rat model of AAV alpha-synucleinopathy. J. Neurosci. Off. J. Soc. Neurosci. 29, 3365–3373 (2009).

50. Tagliaferro, P. et al. An early axonopathy in a hLRRK2(R1441G) transgenic model of Parkinson disease. Neurobiol. Dis. 82, 359–371 (2015).

51. Zarkali, A., Thomas, G. E. C., Zetterberg, H. & Weil, R. S. Neuroimaging and fluid biomarkers in Parkinson’s disease in an era of targeted interventions. Nat. Commun. 15, 5661 (2024).

52. Huang, L.-C. et al. Functional connectivity compensation in Parkinson’s disease with freezing of gait. Eur. J. Neurosci. 60, 6279–6290 (2024).

53. Johansson, M. E., Toni, I., Kessels, R. P. C., Bloem, B. R. & Helmich, R. C. Clinical severity in Parkinson’s disease is determined by decline in cortical compensation. Brain 147, 871–886 (2024).

54. Johansson, M. E., Toni, I., Bloem, B. R. & Helmich, R. C. Parkinson’s disease progression is shaped by longitudinal changes in cerebral compensation. 2025.03.21.25324393 Preprint at 10.1101/2025.03.21.25324393 (2025).

55. Arnatkeviciute, A., Fulcher, B. D., Bellgrove, M. A. & Fornito, A. Imaging Transcriptomics of Brain Disorders. Biol. Psychiatry Glob. Open Sci. 2, 319–331 (2022).

56. Arnatkeviciute, A., Markello, R. D., Fulcher, B. D., Misic, B. & Fornito, A. Toward Best Practices for Imaging Transcriptomics of the Human Brain. Biol. Psychiatry 93, 391–404 (2023).

57. Postuma, R. B. et al. MDS clinical diagnostic criteria for Parkinson’s disease. Mov. Disord. 30, 1591–1601 (2015).

58. Litvan, I. et al. Diagnostic Criteria for Mild Cognitive Impairment in Parkinson’s Disease: Movement Disorder Society Task Force Guidelines. Mov. Disord. Off. J. Mov. Disord. Soc. 27, 349–356 (2012).

59. Creavin, S. T. et al. Mini-Mental State Examination (MMSE) for the detection of dementia in clinically unevaluated people aged 65 and over in community and primary care populations. Cochrane Database Syst. Rev. (2016) doi:10.1002/14651858.CD011145.pub2.

60. Dalrymple-Alford, J. C. et al. The MoCA: well-suited screen for cognitive impairment in Parkinson disease. Neurology 75, 1717–1725 (2010).

61. Stroop, J. R. Studies of interference in serial verbal reactions. J. Exp. Psychol. 18, 643–662 (1935).

62. Wechsler Memory Scale | Fourth Edition. https://www.pearsonassessments.com/en-us/Store/Professional-Assessments/Cognition-%26-Neuro/Wechsler-Memory-Scale-%7C-Fourth-Edition/p/100000281.

63. Rende, B., Ramsberger, G. & Miyake, A. Commonalities and differences in the working memory components underlying letter and category fluency tasks: a dual-task investigation. Neuropsychology 16, 309–321 (2002).

64. Warrington, E. K. The Graded Naming Test: A Restandardisation. Neuropsychol. Rehabil. 7, 143–146 (1997).

65. Warrington, E. K. Recognition Memory Test: Manual. (UKNFER-Nelson, Berkshire, 1984).

66. Hooper, H. Hooper Visual Organization Test (VOT) Manual. (CA: Western Psychological Services, Los Angeles, 1983).

67. Benton, A. L., Varney, N. R. & Hamsher, K. deS. Visuospatial Judgment: A Clinical Test. Arch. Neurol. 35, 364–367 (1978).

68. Goetz, C. G. et al. Movement Disorder Society-sponsored revision of the Unified Parkinson’s Disease Rating Scale (MDS-UPDRS): scale presentation and clinimetric testing results. Mov. Disord. Off. J. Mov. Disord. Soc. 23, 2129–2170 (2008).

69. Shumway-Cook, A., Brauer, S. & Woollacott, M. Predicting the Probability for Falls in Community-Dwelling Older Adults Using the Timed Up & Go Test. Phys. Ther. 80, 896– 903 (2000).

70. Walker, M. P. et al. The Clinician Assessment of Fluctuation and the One Day Fluctuation Assessment Scale. Two methods to assess fluctuating confusion in dementia. Br. J. Psychiatry J. Ment. Sci. 177, 252–256 (2000).

71. Lee, D. R. et al. The dementia cognitive fluctuation scale, a new psychometric test for clinicians to identify cognitive fluctuations in people with dementia. Am. J. Geriatr. Psychiatry Off. J. Am. Assoc. Geriatr. Psychiatry 22, 926–935 (2014).

72. Bjelland, I., Dahl, A. A., Haug, T. T. & Neckelmann, D. The validity of the Hospital Anxiety and Depression Scale. An updated literature review. J. Psychosom. Res. 52, 69– 77 (2002).

73. Pfeffer, R. I., Kurosaki, T. T., Harrah, C. H., Chance, J. M. & Filos, S. Measurement of functional activities in older adults in the community. J. Gerontol. 37, 323–329 (1982).

74. Sletten, D. M., Suarez, G. A., Low, P. A., Mandrekar, J. & Singer, W. COMPASS 31: a refined and abbreviated Composite Autonomic Symptom Score. Mayo Clin. Proc. 87, 1196–1201 (2012).

75. Stiasny-Kolster, K. et al. The REM sleep behavior disorder screening questionnaire-- a new diagnostic instrument. Mov. Disord. Off. J. Mov. Disord. Soc. 22, 2386–2393 (2007).

76. Papapetropoulos, S. et al. A questionnaire-based (UM-PDHQ) study of hallucinations in Parkinson’s disease. BMC Neurol. 8, 21 (2008).

77. Tomlinson, C. L. et al. Systematic review of levodopa dose equivalency reporting in Parkinson’s disease. Mov. Disord. 25, 2649–2653 (2010).

78. Vaculčiaková, L. et al. Combining navigator and optical prospective motion correction for high-quality 500 μm resolution quantitative multi-parameter mapping at 7T. Magn. Reson. Med. 88, 787–801 (2022).

79. Callaghan, M. F. et al. Example dataset for the hMRI toolbox. Data Brief 25, 104132 (2019).

80. Corbin, N., Acosta-Cabronero, J., Malik, S. J. & Callaghan, M. F. Robust 3D Bloch-Siegert based B1+ mapping using multi-echo general linear modeling. Magn. Reson. Med. 82, 2003–2015 (2019).

81. Balbastre, Y. et al. Correcting inter-scan motion artifacts in quantitative R1 mapping at 7T. Magn. Reson. Med. 88, 280–291 (2022).

82. Schaefer, A. et al. Local-Global Parcellation of the Human Cerebral Cortex from Intrinsic Functional Connectivity MRI. Cereb. Cortex N. Y. N 1991 28, 3095–3114 (2018).

83. Vos de Wael, R., et al. BrainSpace: a toolbox for the analysis of macroscale gradients in neuroimaging and connectomics datasets. *Commun*. Biol. 3, 103 (2020).

84. Coifman, R. R. et al. Geometric diffusions as a tool for harmonic analysis and structure definition of data: diffusion maps. Proc. Natl. Acad. Sci. U. S. A. 102, 7426–7431 (2005).

85. Langs, G., Golland, P. & Ghosh, S. S. Predicting Activation Across Individuals with Resting-State Functional Connectivity Based Multi-Atlas Label Fusion. Med. Image Comput. Comput.-Assist. Interv. MICCAI Int. Conf. Med. Image Comput. Comput.-Assist. Interv. 9350, 313–320 (2015).

86. Worsley, K. et al. SurfStat: A Matlab toolbox for the statistical analysis of univariate and multivariate surface and volumetric data using linear mixed effects models and random field theory. NeuroImage 47, S102 (2009).

87. Markello, R. D. et al. neuromaps: structural and functional interpretation of brain maps. Nat. Methods 19, 1472–1479 (2022).

88. Arnatkevic Iūtė, A., Fulcher, B. D. & Fornito, A. A practical guide to linking brain-wide gene expression and neuroimaging data. NeuroImage 189, 353–367 (2019).

89. Markello, R., Shafiei, G., Zheng, Y.-Q. & Mišić, B. abagen: A toolbox for the Allen Brain Atlas genetics data. Zenodo (2020) 10.5281/zenodo.3726257.

90. Arloth, J., Bader, D. M., Röh, S. & Altmann, A. Re-Annotator: Annotation Pipeline for Microarray Probe Sequences. PLOS ONE 10, e0139516 (2015).

91. Amunts, K. et al. BigBrain: An Ultrahigh-Resolution 3D Human Brain Model. Science 340, 1472–1475 (2013).

92. Wagstyl, K. et al. Mapping Cortical Laminar Structure in the 3D BigBrain. Cereb. Cortex 28, 2551–2562 (2018).

93. Paquola, C. et al. The BigBrainWarp toolbox for integration of BigBrain 3D histology with multimodal neuroimaging. eLife 10, e70119 (2021).

94. Blauwendraat, C., Nalls, M. A. & Singleton, A. B. The genetic architecture of Parkinson’s disease. Lancet Neurol. 19, 170–178 (2020).

95. Alexander-Bloch, A. et al. On testing for spatial correspondence between maps of human brain structure and function. NeuroImage 178, 540–551 (2018).

96. Zarkali, A. et al. Dementia risk in Parkinson’s disease is associated with interhemispheric connectivity loss and determined by regional gene expression. NeuroImage Clin. 28, 102470 (2020).

97. Thomas, George et al. Regional brain iron and gene expression provide insights into neurodegeneration in Parkinson’s disease. Brain (2021).

98. Altmann, A. et al. Analysis of brain atrophy and local gene expression in genetic frontotemporal dementia. Brain Commun. doi:10.1093/braincomms/fcaa122.

99. Romero-Garcia, R., Warrier, V., Bullmore, E. T., Baron-Cohen, S. & Bethlehem, R. A. I. Synaptic and transcriptionally downregulated genes are associated with cortical thickness differences in autism. Mol. Psychiatry 24, 1053–1064 (2019).

100. Raudvere, U. et al. g:Profiler: a web server for functional enrichment analysis and conversions of gene lists (2019 update). Nucleic Acids Res. 47, W191–W198 (2019).

101. Supek, F., Bošnjak, M., Škunca, N. & Šmuc, T. REVIGO Summarizes and Visualizes Long Lists of Gene Ontology Terms. PLOS ONE 6, e21800 (2011).

102. Tabelow, K. et al. hMRI - A toolbox for quantitative MRI in neuroscience and clinical research. NeuroImage 194, 191–210 (2019).

103. Fulcher, B. D., Arnatkeviciute, A. & Fornito, A. Overcoming false-positive gene-category enrichment in the analysis of spatially resolved transcriptomic brain atlas data. Nat. Commun. 12, 2669 (2021).

